# On-target toxicity of GSPT1 molecular glue degraders in mice

**DOI:** 10.64898/2026.02.14.705470

**Authors:** Yi-Sheng Pu, Yi-Min He, Xiao-Fan Wei, Yu Zhang, Jie-Fu Fang, Bo Peng, Yong Cang

**Affiliations:** School of Life Science and Technology, ShanghaiTech University, 393 Middle Huaxia Road, Shanghai 201210, China

**Keywords:** humanized CRBN, degradation-resistant GSPT1, molecular glue degraders, animal models for drug safety

## Abstract

Thalidomide is teratogenic in humans but not in rodents due to species-specific differences in the sequence of Cereblon (CRBN), an E3 ubiquitin ligase targeted by thalidomide and its derivative molecular glue degraders (MGDs). This species divergence has hindered the accurate prediction of MGD-induced toxicities in standard laboratory animals. GSPT1 MGDs, such as CC-90009, have shown potent anticancer activities in preclinical models and leukemia patients; however, their clinical development was challenging due to severe adverse effects. This highlights the critical need to characterize on-target toxicities in relevant animal models to exploit the therapeutic safety of this class of MGDs. Here, we generated humanized *Crbn*^V380E^ and *Crbn*^V380E/I391V^ knock-in mouse strains, in combination with a degradation-resistant *Gspt1*^G574N^ strain, to interrogate the *in vivo* on-target effects of CC-90009 and its analog CC-885. We found that targeted GSPT1 depletion in mice led to rapid mortality, preceded by multiple dysfunctions including intestinal obstruction, liver damage, splenic atrophy, and hematological abnormalities. Remarkably, these toxicities, along with the underlying transcriptional perturbations, were completely rescued by the undegradable *Gspt1*^G574N^ mutant, establishing a definitive causal link between GSPT1 degradation and systemic injury. Induced proximity and degradation proteomic analyses revealed that GSPT1 loss triggered a secondary downregulation of many proteins, including MYC, PLK1 and CDK4, which were not directly recruited by these MGDs to CRBN. Collectively, our data define the *in vivo* on-target toxicities associated with endogenous GSPT1 degradation and provide a genetic framework to guide the preclinical safety evaluation of CRBN-based MGD therapeutics.

## Introduction

The development of CRL4^CRBN^-based molecular glue degraders (MGDs) and proteolysis-targeting chimeras (PROTACs) has revolutionized oncology^1,2^ by enabling the degradation of previously considered “undruggable” proteins^3–6^. A significant breakthrough in this field is the targeted degradation of GSPT1 (G1 to S phase transition 1), which, unlike zinc-finger proteins, lacks the characteristic C2H2 domain. GSPT1, also called eRF3a, is a translation termination factor that is ubiquitously expressed and essential for maintaining cellular proteostasis^7^. CC-885 was the first GSPT1 MGD^8^ to be identified, and CC-90009 is its more selective clinical derivative^9,10^; these molecules facilitate the recruitment of GSPT1 to the CRBN E3 ligase for proteasomal degradation. The resulting depletion of GSPT1 triggers extensive translational readthrough^11–13^, leading to robust apoptosis in malignant cells and offering substantial therapeutic potential across diverse malignancies, including relapsed/refractory AML, MYC-driven tumors and metastatic castration-resistant prostate cancer^14–16^.

Despite this therapeutic potential, the clinical development of GSPT1 degraders has encountered several significant setbacks due to severe systemic toxicities. While CC-885 is restricted by off-target co-degradation of proteins like PLK1^17^, CDK4^18^ and BNIP3L^19^, the more selective CC-90009 was recently discontinued from clinical trials (NCT03363373) due to dose-limiting adverse events^10^. Other candidates, such as MRT-2359 (NCT05546268) and GSPT1 degrader-antibody conjugates, ORM-5029 (NCT05521230), have also encountered similar safety hurdles. Despite these setbacks, the pipeline continues to expand with next-generation molecules such as SJ6986 and TD-522^20–22^ and dual-target PROTACs (e.g., WB-156, GBD-9)^22,23^ , underscoring the urgent need to delineate the safety boundaries of GSPT1 degradation. The etiology of GSPT1-mediated toxicity remains a subject of intense debate, with theories ranging from cytokine release syndrome^24,25^ to the secondary depletion of survival factors such as MYC^26–28^. However, resolving this controversy has been hindered by the lack of physiologically relevant models, as wild-type rodents are resistant to CRBN-mediated degradation.

To definitively distinguish GSPT1-specific on-target effects from off-target liabilities *in vivo*, we generated humanized *Crbn*^V380E^ (*Crbn^V/V^*) and *Crbn*^V380E/I391V^ (*Crbn^VI/VI^*) knock-in strains to evaluate CC-885 and CC-90009. By integrating them with a non-degradable *Gspt1*^G574N^ strain^8,13^, we provide the first comprehensive *in vivo* characterization of GSPT1 loss. This study establishes a definitive safety framework for GSPT1-targeting modalities and will enable the development of next-generation degraders to proceed with a clearer understanding of their toxicological limits.

## Results

### Humanized *Crbn* mice recapitulate GSPT1 degradation and drug-induced lethality

Based on previou<u>s</u> findings in cell lines demonstrating that specific CRBN residues dictate sensitivity to GSPT1 degraders^29^, we employed CRISPR/Cas9 technology to generate three humanized knock-in mouse lines: *Crbn*^V380E^, *Crbn*^I391V^, and *Crbn*^V380E/I391V^ (Figure 1A). These newly established strains (*Crbn*^V380E^ and *Crbn*^V380E/I391V^) were developmentally normal, displaying no discernible morphological abnormalities and maintaining normal fertility. Furthermore, the offspring followed a classical Mendelian inheritance pattern, confirming the stable germline transmission of the humanized alleles (Figure S1A).

**Figure 1.**
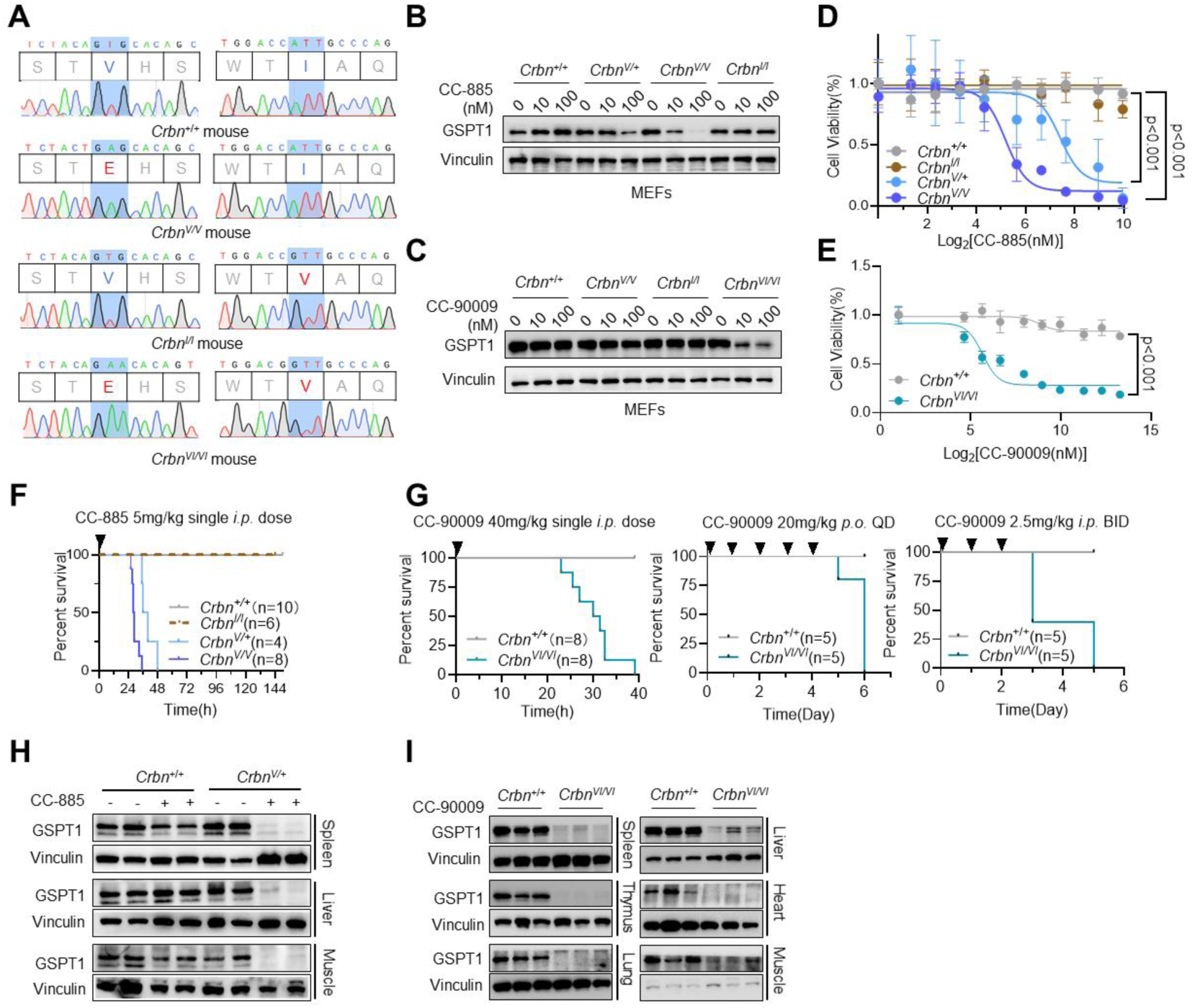
GSPT1 degraders induce lethality in mice following systemic GSPT1 depletion. (A) Validation of *Crbn* mutations by Sanger sequencing. Representative chromatograms displaying the targeted sequences at the 380th (blue highlight, left) and 391st (blue highlight, right) codons of the murine *Crbn* gene. (B and C) Assessment of GSPT1 degradation in *Crbn* mutant MEFs. Immunoblots showing GSPT1 protein levels in MEFs expressing indicated *Crbn* mutants following treatment with (B) CC-885 for 12 h or (C) CC-90009 for 24 h. (D and E) Impact of GSPT1 degradation on cell viability. Survival of *Crbn* mutant MEFs was assessed by CCK-8 assay after 2 days of treatment with (D) CC-885 or (E) CC-90009. Data are presented as mean ±SEM (*n*≥6 biological replicates). *P* values were determined by multiple t-tests. (F and G) Survival analysis of *Crbn* mutant mice. Survival curves of mice expressing indicated *Crbn* mutants following administration of (F) CC-885 (5 mg/kg, single *i.p.* dose) or (G) CC-90009 at various doses and routes (40 mg/kg, single *i.p*.; 20 mg/kg, *p.o.*, QD; or 2.5 mg/kg, *i.p.*, BID). (H and I) Tissue-specific GSPT1 depletion in vivo. Immunoblots of endogenous GSPT1 levels in the indicated organs collected from mice treated with (H) 5 mg/kg CC-885 (*i.p.*, 40 h) or (I) 40 mg/kg CC-90009 (*i.p.*, 30 h).

With these models established, we sought to validate their capacity to support drug-induced degradation of endogenous GSPT1. We performed a systematic evaluation using the molecular glues CC-885 and CC-90009 across various tissues.

Treatment of mouse embryonic fibroblasts (MEFs) derived from these models *in vitro* revealed that CC-885 selectively degraded GSPT1 in *Crbn^V/V^* MEFs (Figure 1B), while CC-90009-mediated degradation was restricted to *Crbn^VI/VI^* MEFs (Figure 1C). Quantitative proteomics further demonstrated that CC-885 triggered rapid and selective degradation of endogenous GSPT1 within 5 h in these engineered *Crbn^V/V^* cells (Figure S1B–S1D). Similarly, CC-90009 was validated to specifically degrade GSPT1 in *Crbn^VI/VI^* cells^13^. Correspondingly, both compounds exhibited potent cytotoxicity only in MEFs undergoing GSPT1 loss (Figure 1D and 1E), whereas wild-type and *Crbn^I/I^* cells remained entirely resistant to both degraders and their associated toxicity.

Administration of GSPT1 degraders *in vivo* resulted in rapid lethality. A single intraperitoneal (*i.p.*) dose of CC-885 (5 mg/kg) led to acute morbidity in *Crbn^V/V^* mice, characterized by decline in body temperature, profound lethargy, and subsequent death, but not in wild-type or *Crbn^I/I^* littermates (Figure 1F). CC-90009 elicited a remarkably similar toxicological profile in *Crbn^VI/VI^* mice across various dosing regimens (Figure 1G). Immunoblot analysis confirmed the selective depletion of endogenous GSPT1 by CC-885 in *Crbn^V/V^* mutants (Figure 1H) and by CC-90009 in *Crbn ^VI/VI^* mutants (Figure 1I) , with no depletion observed in controls.

Collectively, these findings demonstrate that the *Crbn^V/V^*and *Crbn^VI/VI^* strains successfully recapitulate the degradation of endogenous GSPT1 by CC-885 and CC-90009, leading to systemic toxicity and lethality.

### GSPT1 degraders induce multi-organ toxicity in humanized *Crbn* mice

To elucidate the cause of the rapid lethality observed in humanized mice, we assessed potential tissue damage. Given the ubiquitous expression of GSPT1, we examined both highly proliferative tissues (intestine and spleen) and relatively quiescent organs (liver).

Intestinal abnormalities were among the most obvious manifestations in CC-885-treated *Crbn^V/V^* mice, characterized by marked gastrointestinal (GI) distension with accumulation of turbid, yellow exudate within the intestinal lumen (Figure 2A). Physiological monitoring revealed hypotension specific to *Crbn^V/V^* mice (Figure S2A), accompanied by lethargy and systemic deterioration consequent to severe GI dysfunction. Gram staining of duodenum-jejunum section confirmed extensive bacterial invasion into the mucosa (Figure S2B), which elicited a systemic inflammatory response characterized by markedly elevated neutrophils and monocytes (Figure S3A and S3B). Assessment of gastrointestinal transit using a charcoal meal confirmed profound GI dysfunction, manifested as complete paralytic ileus in treated mice (Figure 2B). We postulated that the observed hypotension was likely secondary to severe ileus, while the breakdown of the intestinal wall and subsequent bacterial translocation served as the primary drivers of systemic toxicity. Consistent with this, histological and immunohistochemical analyses 10 h post-treatment revealed crypt abscesses, villus disruption, and widespread crypt cell apoptosis closely associated with GSPT1 degradation (Figure 2C).

**Figure 2.**
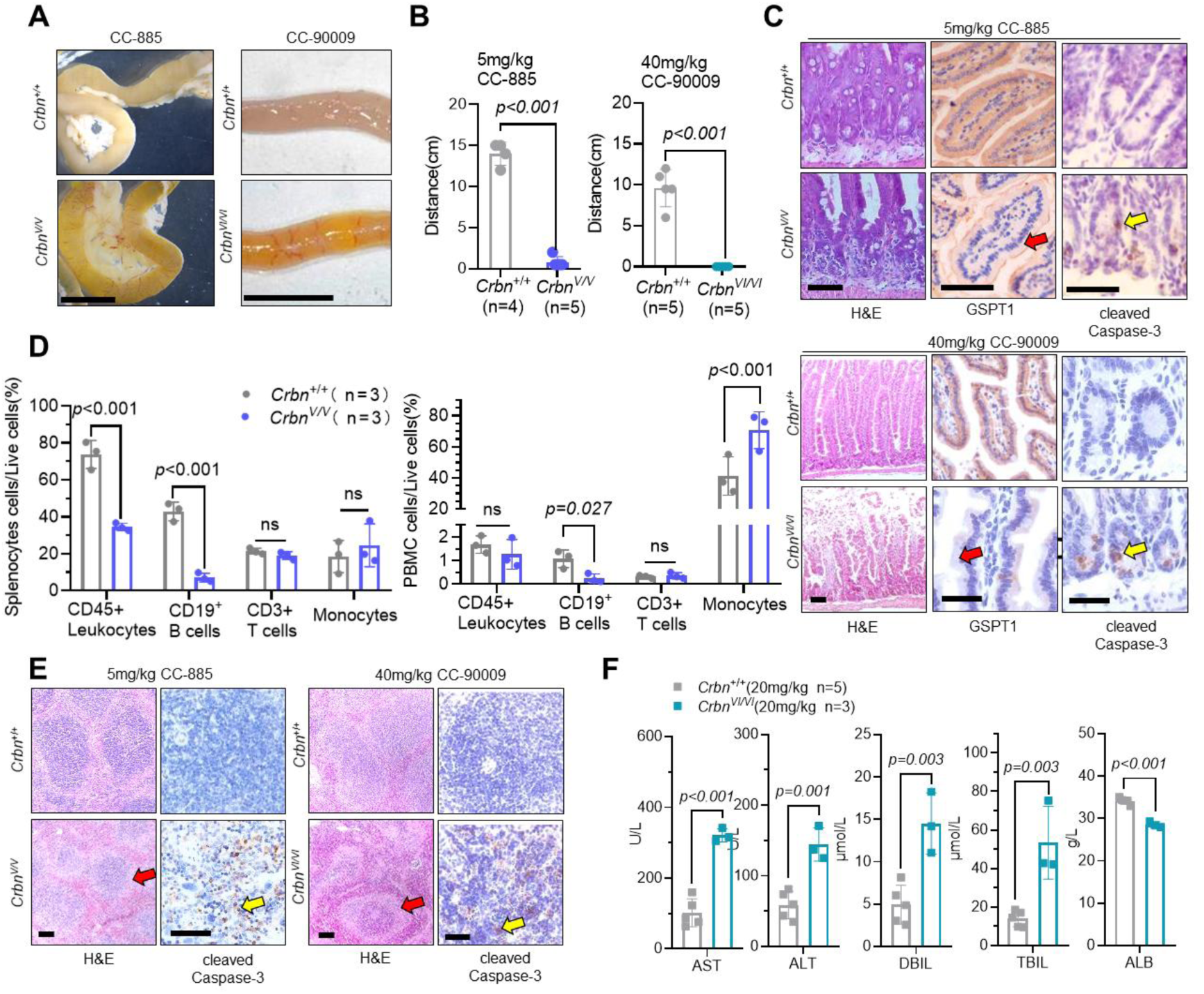
GSPT1 degraders induce proliferating cell apoptosis and multi-organ damage. (A) Intestinal macro-morphology. Stereo microscopic images of duodenum and jejunum from *Crbn*^+/+^ and *Crbn*^V/V^ mice 20 h post-CC-885 (5 mg/kg, *i.p.*). Scale bars, 10 mm. (B) Assessment of gastrointestinal motility. Intestinal transit (charcoal propulsion) in WT or humanized *Crbn* mice 15 h post-CC-885 (5 mg/kg, single *i.p.* dose; left) or CC-90009 (40 mg/kg, single *i.p.* dose; right). (C) Jejunal histology and IHC. H&E and IHC staining for GSPT1 and cleaved Caspase-3 in jejunal sections 20 h (CC-885, 5 mg/kg *i.p.*) or 10 h (CC-90009, 40 mg/kg *i.p.*) post-treatment. Red arrows, GSPT1 depletion; yellow arrows, apoptotic crypt cells. All scale bars, 50μm. (D) Immunophenotyping of hematopoietic populations. Relative counts of monocytes, CD3^+^ T cells, and CD19^+^ B cells in splenocytes and PBMCs from *Crbn*^+/+^ and *Crbn*^V/V^ mice, analyzed by flow cytometry 20 h after a single 5 mg/kg *i.p.* dose of CC-885. (E) Splenic histopathology. H&E and IHC staining for cleaved Caspase-3 in splenic sections following treatment with 5 mg/kg CC-885 (left, 20 h) or 40 mg/kg CC-90009 (right, 10 h). Red arrows, white pulp atrophy; yellow arrows, apoptotic cells. All scale bars, 50μm. (F) Hepatic function. Plasma liver function biomarkers in *Crbn*^+/+^ and *Crbn*^VI/VI^ mice following repeated dosing of CC-90009 (20 mg/kg, *p.o.*, QD). Data: mean ± SD (**B, D, F**); *n*, biological replicates. *P* values: multiple t-tests. All experiments performed ≥3 times with representative data shown. n.s., not significant.

Hematological profiles mirrored those of systemic bacterial infection, characterized notably by lymphopenia. Flow cytometric analysis of peripheral blood mononuclear cells (PBMCs) and splenocytes revealed a significant reduction in CD45^+^ leukocytes, which may be attributed to severe CD19^+^ B cell loss, while CD3^+^ T cells remained largely unaffected (Figure 2D). Histological examination of the spleen revealed extensive apoptosis and white pulp atrophy following treatment (Figure 2E), confirming GSPT1-mediated B-cell toxicity.

Treated humanized mice also exhibited hepatic injury, evidenced by icteric liver or yellowish discoloration of the liver parenchyma (Figure S4A and S4B). While liver weight remained comparable to controls (Figure S4C), plasma biochemistry revealed significant alterations in transaminase levels (ALT, AST), along with shifted bilirubin (DBIL, TBIL) and albumin (ALB) levels, confirming hepatic injury and impaired function (Figure 2F and S4D). Histopathological evaluation also identified hepatic congestion and stasis following drug administration (Figure S4E).

Together, these results demonstrate that GSPT1 degradation in humanized *Crbn* mice elicits severe multi-organ toxicity, particularly within metabolically active and rapidly proliferating tissues, which leads to the observed physiological decline and lethality.

### Degrader-induced lethality is exclusively mediated by on-target GSPT1 depletion

The observation that severe physiological decline and multi-organ failure were consistently coincided with GSPT1 clearance prompted us to investigate whether GSPT1 degradation is the definitive cause of the observed toxicity. Since thalidomide analogues like Pomalidomide (Poma) and GSPT1 degraders share the same binding pocket on CRBN^8^, we initially performed competition assays to evaluate the dependency of toxicity on substrate degradation. Poma effectively outcompeted CC-885 for CRBN binding, thereby inhibiting CC-885-induced degradation (Figure S5A). This pharmacological intervention not only rescued MEFs from cytotoxicity (Figure S5B) but also mitigated lethality in humanized mice (Figure S5C), indicating that the toxicity is primarily mediated by CRBN-dependent substrate degradation rather than off-target effects.

To genetically dissect the role of GSPT1 loss, we generated a non-degradable mouse model. Given the G-loop sequence conservation between human and murine GSPT1, we introduced a glycine-to-asparagine substitution (G574N) into the murine G-loop (Figure S6A), which conferred resistance to the degradation and cytotoxicity effects of both CC-885 and CC-90009 in mouse cell lines (Figure S6B -S6E). We subsequently generated *Gspt1*^G574N^ knock-in mice (*Gspt1^G/G^*) (Figure 3A) and crossed them with humanized *Crbn* strains to generate double-mutant mice which were viable and followed Mendelian inheritance (Figure S6F and S6G). Functional validation in MEFs confirmed that *Gspt1*^G574N^ mutation conferred robust resistance to drug-induced degradation and subsequent apoptosis in humanized *Crbn* cells (Figure 3B and 3C).

**Figure 3.**
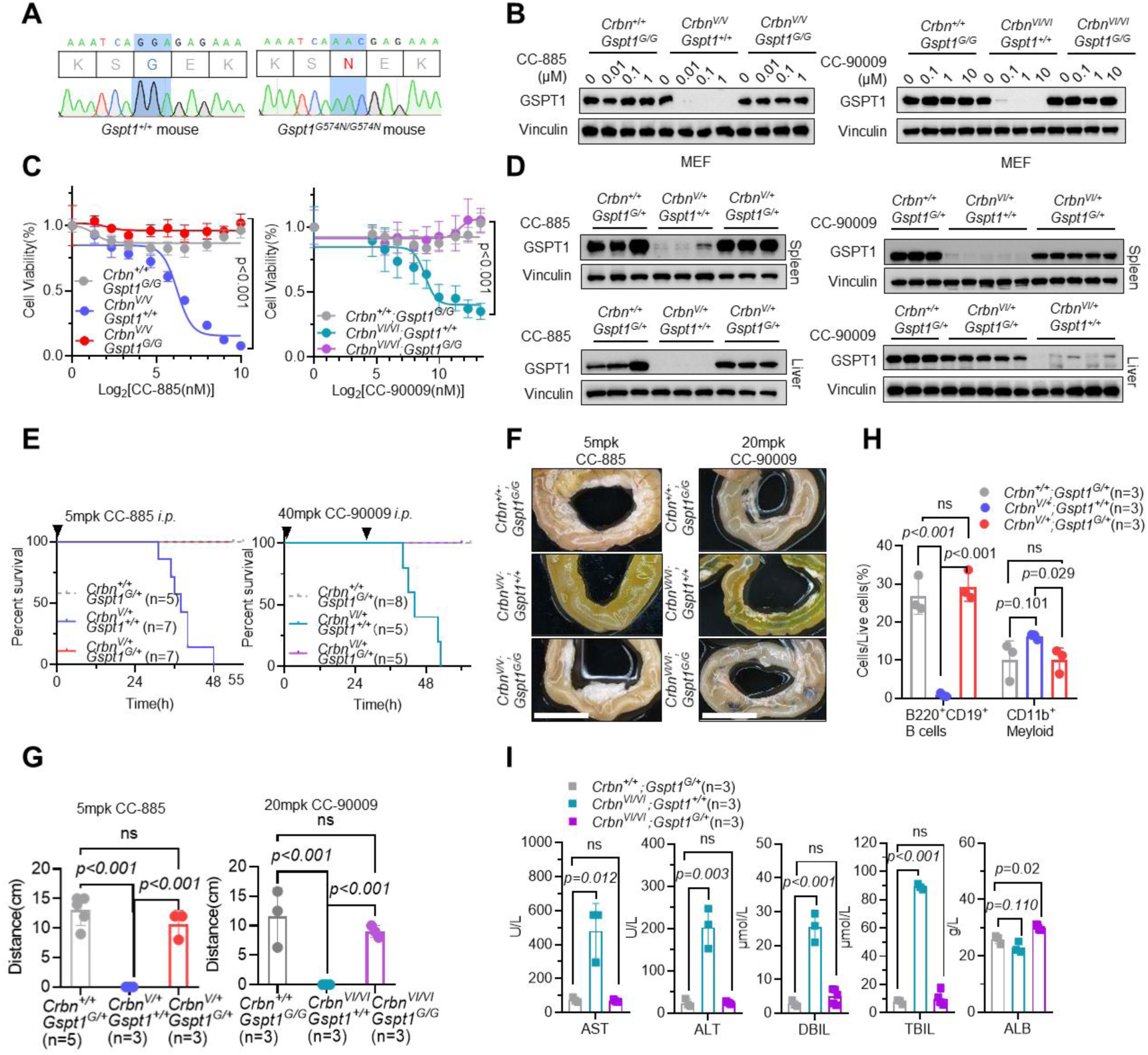
Rescue of GSPT1 degradation prevents lethality and multi-organ damage. (A) Generation of degrader-resistant mice. Sanger sequencing chromatogram verifying the *Gspt1*^G574N^ knock-in mutation. The blue box indicates the codon substitution from GGA (Gly) to AAC (Asn) at position 574. (B) Resistant MEFs. Immunoblot analysis of GSPT1 levels in MEFs of indicated genotypes following treatment with CC-885 (12 h) or CC-90009 (24 h). (C) Cell viability. Viability of MEFs of indicated genotypes after 48 h of continuous treatment with CC-885 (left) or CC-90009 (right). (D) In vivo GSPT1 levels. Western blot analysis of GSPT1 in spleen and liver sections from indicated mouse genotypes after a single *i.p.* injection of CC-885 (5 mg/kg, left) or CC-90009 (40 mg/kg, right). (E) Survival analysis. Survival curves of indicated mouse genotypes following a single *i.p.* dose of 5 mg/kg CC-885 (left) or two *i.p.* doses of 40 mg/kg CC-90009 (right). (F) Intestinal macro-morphology. Representative images of the duodenum and jejunum (top) and corresponding intestinal propulsion (bottom) after CC-885 (5 mg/kg *i.p.*, single dose) or CC-90009 (20 mg/kg *i.p.*, QD for 3 days). Scale bars, 10 mm. (G) GI motility. Intestinal transit measured via charcoal propulsion 15 h post-administration of CC- 885 (5 mg/kg *i.p.*) or CC-90009 (40 mg/kg *i.p.*). (H) Bone marrow immunophenotyping. Flow cytometric quantification of myeloid and B-cell populations in the bone marrow 20 h after a single 5 mg/kg *i.p.* dose of CC-885. (I) Hepatic function. Plasma liver biomarkers in indicated mouse genotypes following repeated CC-90009 dosing (20 mg/kg, *p.o.*, QD for 3 days). Data: mean ±SD (**C, G, H, I**); *n*, biological replicates. *P* values: multiple t-tests. All experiments performed ≥3 times with representative data shown. n.s., not significant.

We next challenged these double mutant mice with previously lethal doses of CC-885 (5 mg/kg) or CC-90009 (40 mg/kg). Neither compound induced GSPT1 degradation in spleen or liver (Figure 3D), indicating that the G574N mutation effectively abrogates systemic degradation of GSPT1 *in vivo.* Crucially, the non-degradable *Gspt1^G574N^* mutation protected humanized *Crbn* mice against the degrader-induced toxicity. These mice remained active and exhibited no signs of lethargy or physical deterioration throughout the observation period (Figure 3E).

The preservation of GSPT1 levels across these diverse tissues conferred complete protection against multi-organ pathology. In both treatment groups, the intestinal morphology of double-mutant mice resembled wild-type controls, with no observable wall thinning, accumulation of turbid fluid, or paralytic ileus (Figure 3F and 3G). Hematological abnormalities were similarly absent in the double-mutant cohort. Furthermore, the maintenance of neutrophil levels (confirmed in bone marrow, Figure 3H), along with the preservation of lymphocytes and platelet counts (Figure S7B), collectively underscore that the systemic inflammatory response was strictly dependent on GSPT1 degradation. Additionally, B-cell populations in the bone marrow were protected, confirming that B-cell loss is a direct consequence of GSPT1 degradation (Figure 3H). Hepatic function was also preserved, as serum levels of ALT, AST, and bilirubin returned to baseline (Figure 3I).

Transcriptomic profiling demonstrated that GSPT1 protection maintained global gene expression in the spleen and liver to wild-type levels (Figure 4A and 4B). Gene set enrichment analysis (GSEA) revealed that the G574N mutation effectively reversed the transcriptional signatures of toxicity. Specifically, in the spleen, GSPT1 protection preserved DNA replication and cell cycle pathways while maintaining normal immune response signatures. In the liver, essential metabolic and complement pathways were similarly preserved, alongside a significant alleviation of oxidative stress (Figure S7C and S7D).

**Figure 4.**
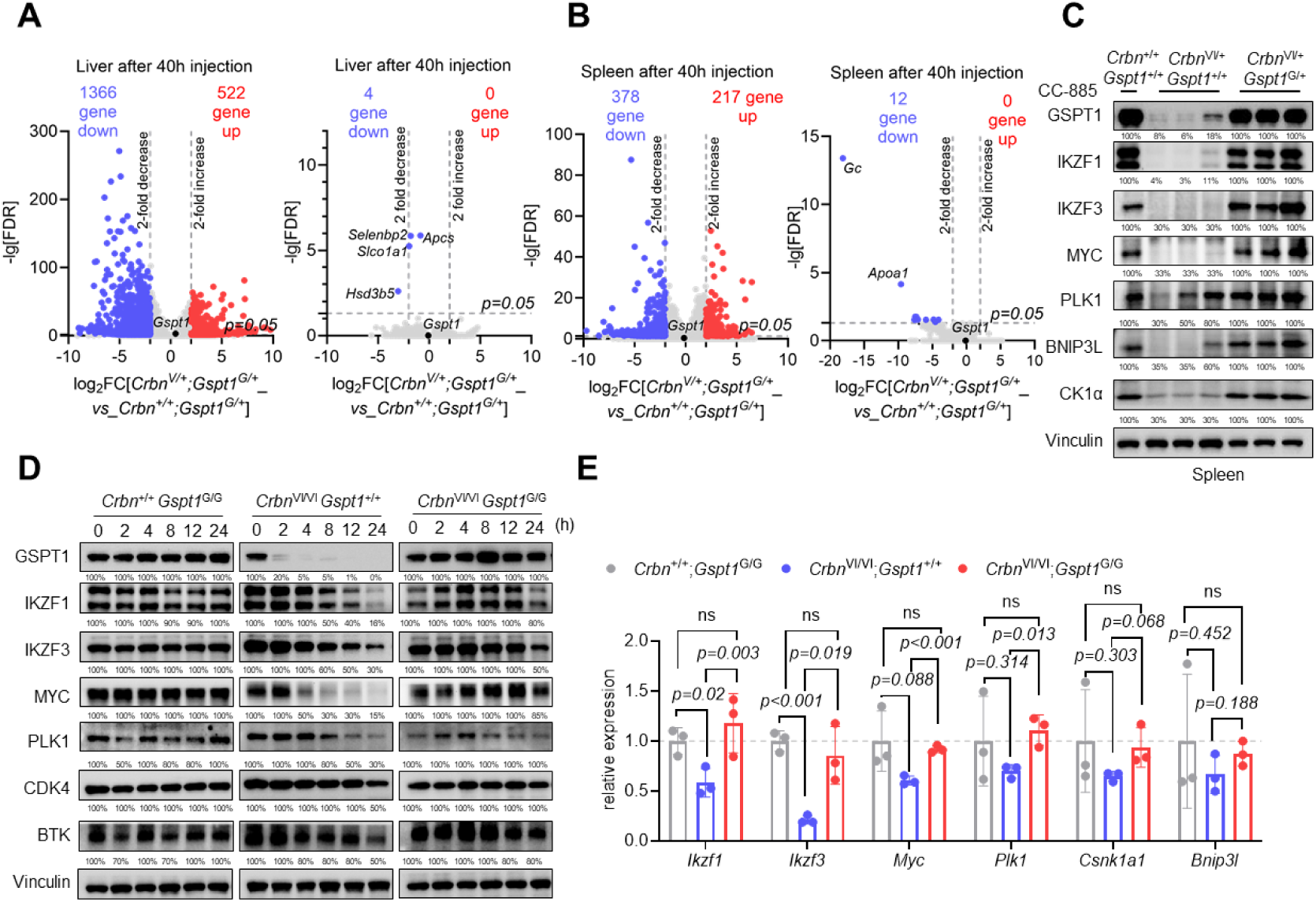
GSPT1 degradation leads to the downregulation of other target proteins. (A and B) Transcriptomic profiling. RNA-seq analysis showing differential gene expression in the liver (A) and spleen (B) of *Crbn*^+/+^; *Gspt1^G/+^*, *Crbn^V/+^*; *Gspt1^+/+^* and *Crbn^V/+^*; *Gspt1^G/+^* mice at 40 h post-injection of 5 mg/kg CC-885 (*i.p.*). (C) Downregulation of GSPT1 and associated proteins. Western blot analysis of GSPT1 and proteins involved in cell cycle and survival, including MYC, PLK1, BNIP3L, and CK1α, in splenic tissues. (D) Dynamic protein expression patterns. Time-course Western blot of GSPT1, MYC, PLK1, CDK4, and BTK levels in splenocytes following CC-885 treatment. Note: In G574N mutant cells, the levels of these proteins were largely sustained, correlating with GSPT1 stability. (E) mRNA levels. qRT-PCR quantification of *Myc*, *Plk1*, *Bnip3l*, and *Csnk1a1* mRNA levels in the spleen, showing the transcriptional response following GSPT1 depletion. Data are presented as mean ±SEM (*n* = 3 biological replicates). *P* values were determined by multiple t-tests.

Collectively, these findings provide definitive genetic evidence that the cascade of intestinal failure, hematopoietic failure, and lethal systemic shock induced by these degraders is driven exclusively by GSPT1 depletion.

### GSPT1 loss drives secondary proteome collapse via transcriptional repression

Transcriptomic profiling revealed that GSPT1 degradation induces a broad downregulation of gene expression, underscoring that downstream regulatory consequences extends beyond the immediate loss of the neo-substrate. While compounds like CC-885 have been reported to directly target substrates such as IKZF1/3^8^, BNIP3L^19^, PLK1^17^, and CDK4^18^, our data showed that in murine tissues, these proteins decreased significantly only upon GSPT1 degradation (Figure 4C).

This dependency was consistently recapitulated in murine splenocytes (Figure 4D) and human cell lines, including MOLT-4 (Figure S8A and S8B) and 293T (Figure S8C). Furthermore, splenic transcriptomic and qRT-PCR analyses confirmed the downregulation of the transcripts encoding these targets (Figure 4E). In sum, GSPT1 depletion instigates a broad transcriptional collapse, which dictates the downstream reduction in protein levels.

To systematically dissect direct versus secondary targets, we performed quantitative proteomic analysis in degradation-resistant *GSPT1^G575N^* mutant 293T cells following CC-885 treatment. This comparative approach allowed us to distinguish between proteins directly recruited by CC-885—such as GSPT2, WIZ, and CHD7—and a wider array of proteins whose abundance decreased only as a consequence of GSPT1 depletion, including TGIF2, EEF2K, MYC, CHEK1, PLK1 and CDK4 (Figure 5A, 5B and S9A). To definitively determine whether these reductions resulted from direct recruitment or downstream effects, we employed a CRBN-TurboID proximity labeling system (Figure 5C, S10A, and S10B). Mass spectrometry analysis confirmed that TGIF2, EEF2K, MYC, CHEK1, PLK1 and CDK4 were not directly recruited to the CRBN complex by the compounds (Figure S10C and S10D). Instead, their depletion represents a downstream proteomic signature driven by the transcriptional collapse following GSPT1 degradation.

**Figure 5.**
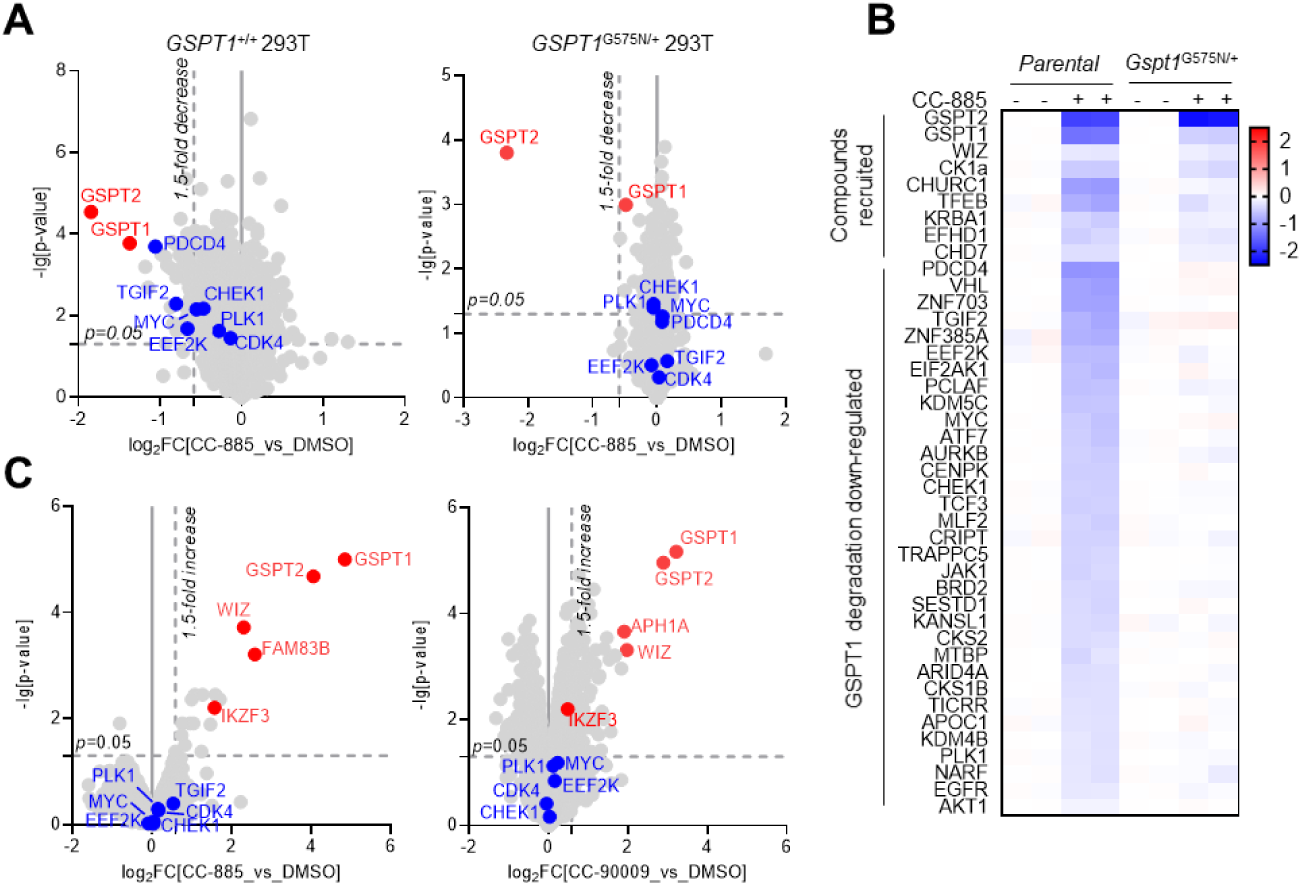
Global proteomic profiling identifies GSPT1 degradation-dependent protein alterations. (A) Quantitative proteomics. Volcano plots showing changes in protein abundance in WT (right) and *GSPT1^G575N^* mutant (left) 293T cells following 24 h treatment with 100 nM CC-885. (B) Classification of GSPT1 degradation-dependent proteins and CC-885 recruited targets. Heatmap showing the relative abundance of selected proteins. These include putative direct CRBN neosubstrates (degraded in both genotypes) and GSPT1-dependent proteins (only downregulated in WT cells upon GSPT1 depletion) (C) CRBN-TurboID proximity labeling. TurboID identifying potential proteins recruited by CRBN in the presence of 1 μM CC-885 (left) or 10 μM CC-90009 (right) in 293T cells.

In conclusion, these findings demonstrate that GSPT1 loss precipitates a widespread secondary collapse of the cellular proteome, primarily driven by a cascade of transcriptional downregulation that extends far beyond direct neo-substrate recruitment. This discovery suggests that the narrow therapeutic window of GSPT1-targeting therapies is intrinsically linked to this broad, secondary disruption of the cellular proteome.

## Discussion

Targeting GSPT1 for degradation has long been recognized for its exceptional anti-tumor potential and promising druggability. However, the clinical trajectory of GSPT1-targeting molecular glues has been fraught with challenges; over the past decade, the majority of these candidates have withdrawn from clinical development due to safety concerns. Despite these setbacks, the precise mechanistic basis of GSPT1-mediated systemic toxicity has remained largely elusive. Consequently, delineating the safety boundaries of GSPT1 degradation is of paramount importance for successful clinical translation, especially as a new generation of GSPT1-targeting degraders continues to enter clinical pipelines.

Our study addresses this critical gap by establishing that the lethal systemic shock observed is a direct, on-target consequence of GSPT1 depletion. By leveraging humanized *Crbn* mice and non-degradable *Gspt1^G574N^* mutants, we provided definitive evidence that the degradation of GSPT1 is the direct and primary cause of lethal multi-organ toxicity—including gastrointestinal injury, hematological abnormalities (such as B-cell depletion), and hepatic impairment. Importantly, this toxicological profile closely mirrors the severe adverse events reported in clinical patients, like hyperbilirubinemia, hypotension, sepsis and thrombocytopenia^30^, confirming that the observed pathology is an on-target, mechanism-driven effect rather than an off-target liability. Therefore, the therapeutic window for this approach is inherently narrow due to profound disruption of essential tissue homeostasis. Future clinical strategies will thus depend not only on identifying tumor types with intrinsic hypersensitivity as suitable patient populations, but also on designing specific drug combination regimens to expand the safety margin.

Beyond the systemic level, our molecular analysis reveals that GSPT1 loss triggers a secondary collapse of the cellular proteome, fundamentally driven by widespread transcriptional downregulation. We demonstrate that the depletion of multiple survival-related proteins and critical cell cycle regulators is not a result of direct neo-substrate recruitment, but a secondary consequence of the transcriptional and translational disruption caused by GSPT1 loss. This phenomenon carries profound implications for the field of targeted protein degradation, highlighting a significant pitfall in proteomics-based “degradomics” where many identified “targets” may, in fact, be indirect downstream effects. Consequently, our work mandates a more rigorous standard for target validation: the use of non-degradable mutants or proximity labeling systems is essential to definitively distinguish true molecular glue substrates from indirect, GSPT1-dependent proteomic alterations.

In light of these findings, our study underscores the indispensable value of humanized *Crbn* models in bridging the gap between preclinical research and clinical outcomes. Species-specific differences in CRBN have long obscured the true toxicological profile of molecular glue degraders, leading to unforeseen risks in human subjects. By accurately recapitulating clinical adverse events, our humanized *Crbn* strains provide a reliable and predictive “gold-standard” platform. We therefore propose that such humanized models should be routinely employed in the preclinical evaluation of next-generation CRBN-based degraders to rigorously assess both their pharmacological efficacy and toxicological safety before clinical entry.

## Limitation

The study employed humanized *Crbn* (*Crbn*^V380E^ *and Crbn*^V380E/I391V^) and *Gspt1*^G574N^ mouse models—resistant to small molecule induced degradation—to demonstrate that pharmacological degradation of endogenous GSPT1 causes extensive multi-organ damage and lethality in mice, indicating that targeted degradation of GSPT1 is associated with substantial systemic toxicity. Future studies delineating the temporal progression of these toxicities will be crucial for identifying potential interventional windows. Furthermore, exploring dose reduction or optimized dosing regimens may help define a viable therapeutic index for this target class, suggesting that while GSPT1-targeting degraders face significant safety challenges, their clinical viability is not entirely precluded. In addition, while our study characterized the general features of other proteins downregulated upon GSPT1 degradation, it lacked further classification and mechanistic insights into these affected proteins.

## Significance

This study was the first to validate the diverse toxic manifestations caused by small molecule-induced degradation of GSPT1 in animal models. These included apoptosis of crypt cells and intestinal obstruction in the lower gastrointestinal tract, bacterial infection accompanied by elevated neutrophil and monocyte counts, a sharp decline in B cell numbers, increased red blood cell counts, hepatotoxicity, and mouse lethality. Furthermore, the study further characterized the profile of proteins that were downregulated upon GSPT1 degradation, including known small molecule recruitable targets such as IKZF1/3, cell cycle–related proteins such as PLK1 and CDK4, as well as other potential targets such as CHEK1 and BTK. Collectively, these findings provide a comprehensive toxicity atlas that links proteomic perturbations to systemic pathologies, offering a critical roadmap for the rational design of next-generation degraders with improved therapeutic windows.

## Acknowledgments

We thank Degron Therapeutics Co. Ltd for data sharing and technical support. We also thank the Multi-Omics Core Facility, Molecular and Cell Biology Core Facility, Molecular Imaging Core Facility, and Animal Core Facility at ShanghaiTech University for mass spectrometry system technical support, electron microscopy system technical support, molecular imaging system technical support, and mouse breeding support. This research was supported by Shanghai Frontiers Science Center for Biomacromolecules and Precision Medicine at ShanghaiTech University. This work was supported by grants from the National Key R&D Program and the National Natural Science Foundation of China (No. 2025YFA1309004).

## Author contributions

Y.S.P. and Y.C. conceived the study; Y.S.P. performed most of the experiments; X.F.W. contributed to the counts of B cells in splenocytes and bone marrows; Y.M.H. contributed to breeding of the mice and data analysis and integration; Y.Z. and J.F.F. provided technical assistance and experimental support; Y.L. and J.Y.C. contributed to the critical revision of the manuscript; Y.S.P., and B.P., wrote the manuscript; Y.C. obtained the funding and supervised the study.

## STAR★Methods

### Key resources table

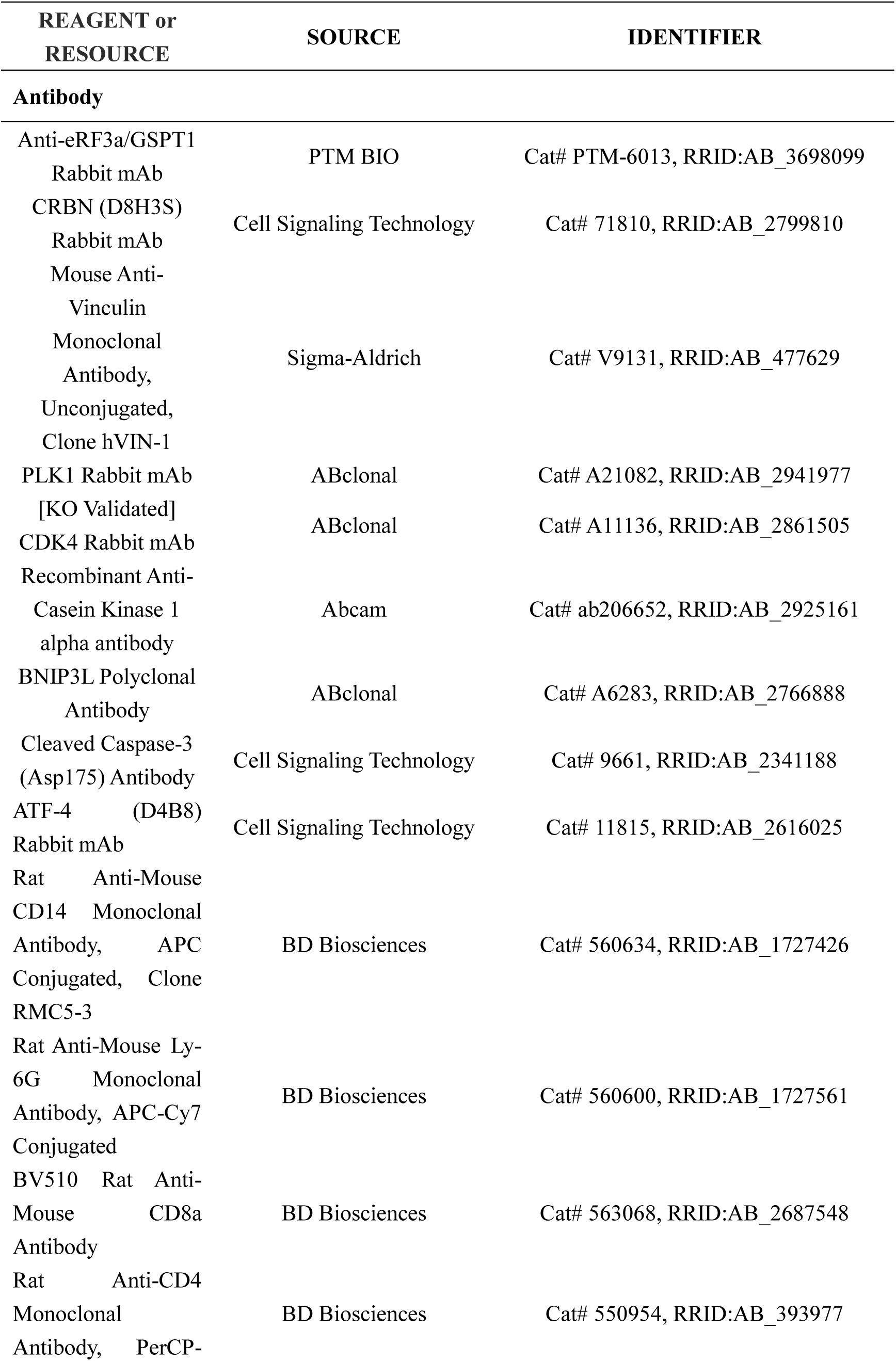
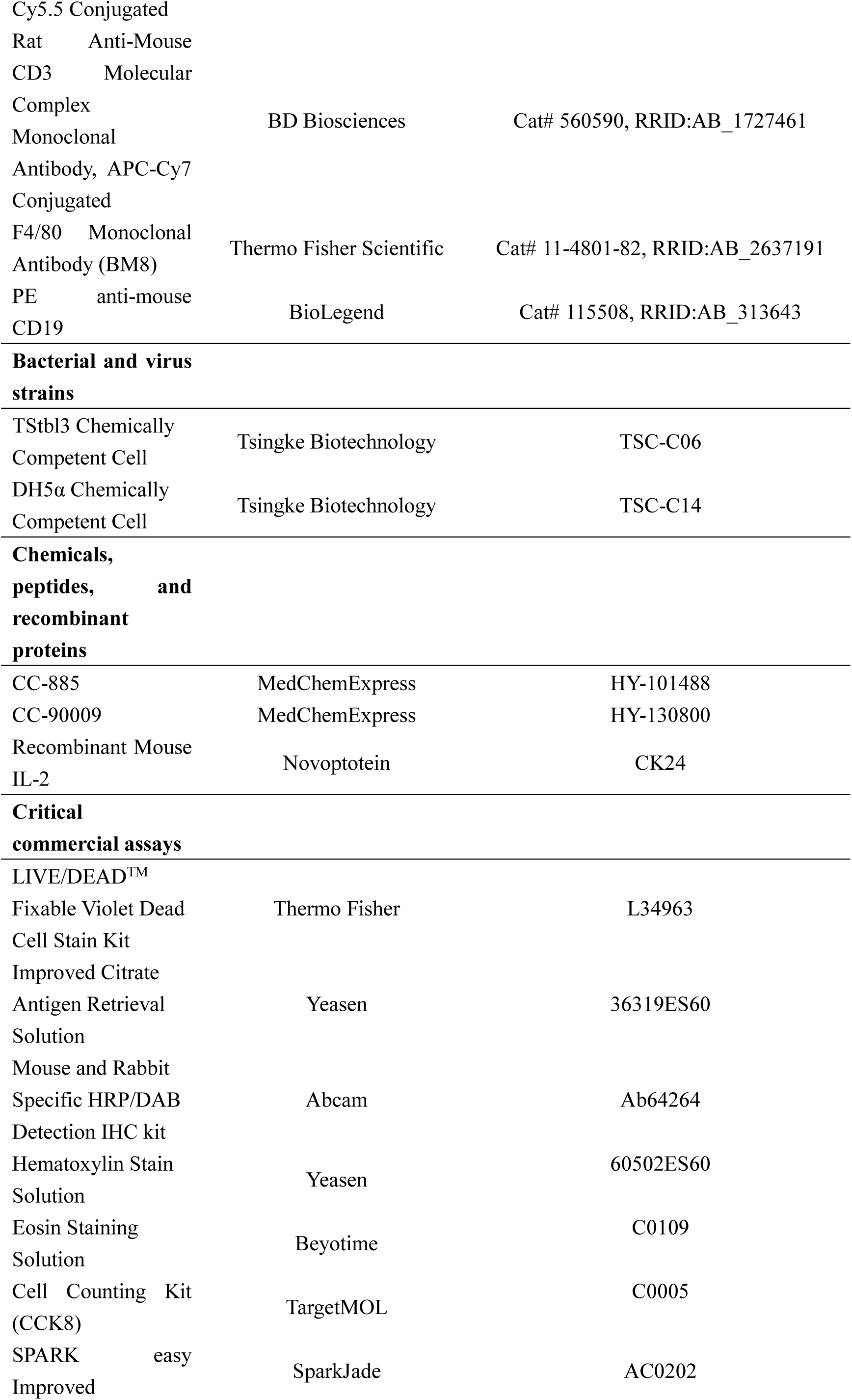
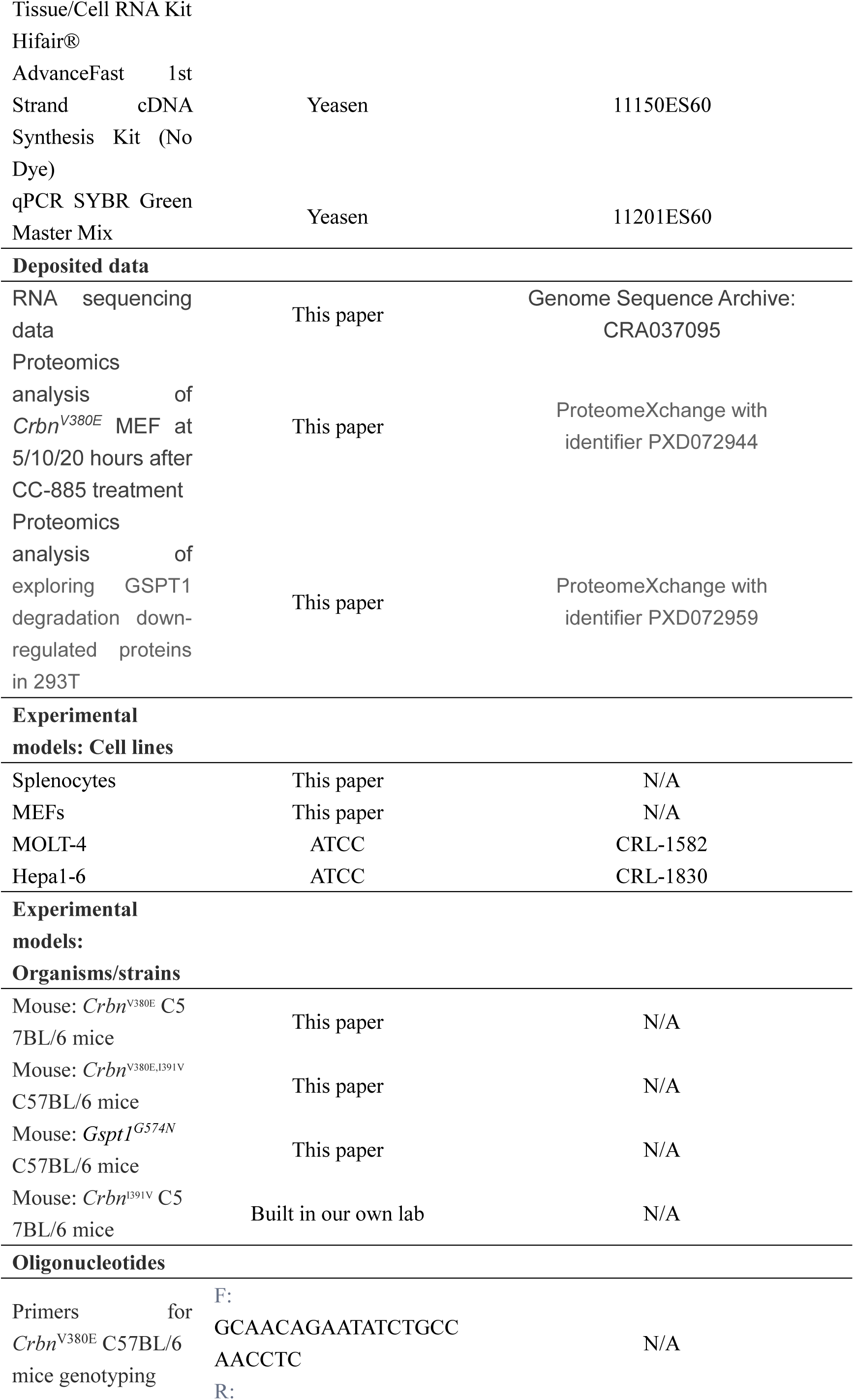
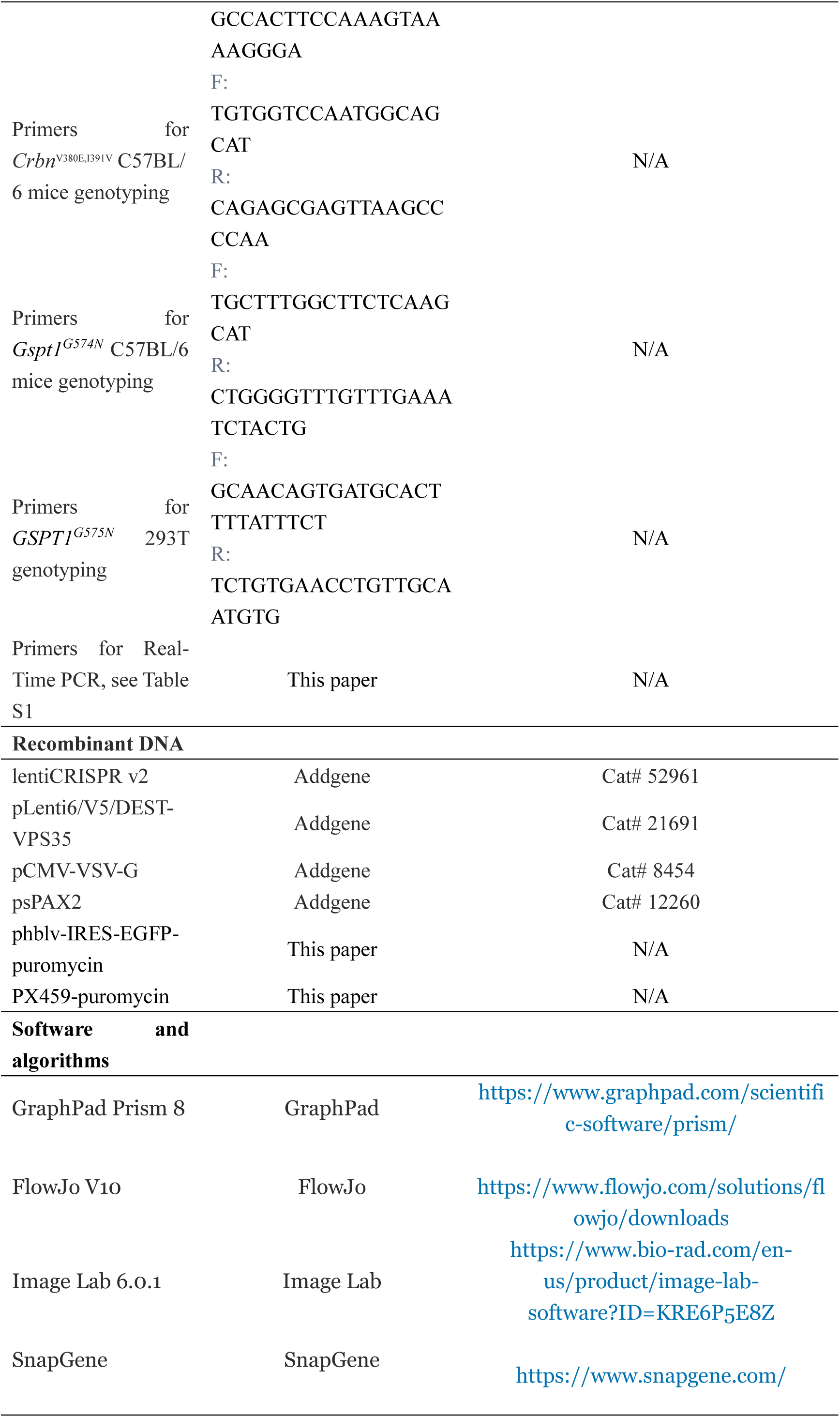

### Resource availability Lead contact

Further information and requests for resources and reagents should be directed to and will be fulfilled by the lead contact, Yong Cang.

### Materials availability

Any plasmids, cell lines, and mouse lines generated in this study are available from the lead contact upon request.

**Data and code availability**

## Method details

### Generation of the *Crbn*^V380E^, *Crbn*^V380E/I391V^ and *Gspt1*^G574N^ mouse model

All mouse strains used in this study were maintained on a C57BL/6 background and generated using the CRISPR-Cas9 system. For the generation of *Crbn^V380E^* mice, a specific single-guide RNA (sgRNA) targeting the murine *Crbn* locus (5’-ACAGTGCACAGCTGGTTTCC-3’) was utilized. Intact sgRNAs were amplified from the PX459 plasmid (Addgene, #48139) using T7 promoter-tagged primers (Forward: 5’-CACCGACAGTGCACAGCTGGTTTCC-3’; Reverse: 3’-AAACCGGAAACCAGCTGTGCACTGTC-5’). For the *Crbn^V380E/I391V^* strain, the same strategy was applied to a pre-existing *Crbn^I391V^* background using an sgRNA targeting the sequence 5’-GGCCTTCTACAGTGCACAGCTGG-3’. The *Gspt1^G574N^*model was generated using an sgRNA targeting exon 14 of the murine *Gspt1* gene (5’-TGCTTGGTAGACAAGAAATC-3’).

To facilitate precise homology-directed repair, specific ssDNA templates were co-injected with the Cas9/sgRNA complex.

- The ssDNA sequence for *Crbn^V380E^*engineering was: 5’-AAAGCGTCCAACCTGAATCTGATAGGCCGGCCTTCTAC**TGA**GCACAGCTG GTTTCCCGGGTAATACAGCTGTTTACTTTTCTTGTTGACTCTTCATTTAGTTTTAGATGAACTTTCTAGGAAGATACAAAACAAACAGGACAGGAATAGTTTGATCACTTCATGAATGGGTTAAAAGCAGGGACATGAGA-3’.
- For *Crbn^V380E/I391V^* mice, the ssDNA template was: 5’-ACTGACTGTGTATAAAGCGTCCAACCTGAATCTGATAGGCCGGCCTTCTA C**AGA**ACACAG**TT**GGTTTCCCGGgtaatacagctgtttacttttct-3’.
- For *Gspt1^G574N^* mice, the ssDNA template was: 5’-GCCTTAATCTGCTTGGTAGACAAGAAATC**AAC**GAGAAAAGTAAGACTCGA CCCCGTTTTGTAAAACAAGATC-3’.

The resulting founders were confirmed by Sanger sequencing and backcrossed to ensure germline transmission. (Note: Targeted mutations are integrated with synonymous mutations to prevent re-cleavage by Cas9 and to facilitate genotyping).

Age-matched C57BL/6 mice were purchased from ShanghaiTech University Animal Core Facility. *Gspt1*^G574N^ mice were bred to *Crbn*^V380E^ or *Crbn*^V380E/I391V^ mice in house. The protocol and any amendment(s) or procedures involving the care and use of animals in this study were in accordance with Shanghai Institutional Animal Care and Use Committee (IACUC) guidelines and approved by IACUC of ShanghaiTech University.

### Cell culture and primary cells extraction

293T, Hepa1-6 cells were maintained in complete DMEM media (Gibco, 10% FBS, and 50 U/ml of Penicillin-Stretomycin). MOLT-4 cells were maintained in complete RPMI 1640 media (Gibco, 10% FBS, and 50 U/ml of Penicillin-Stretomycin). All cell lines used in this study were tested as Mycoplasma-negative using the Universal Mycoplasma Detection Kit (ATCC, 30-1012K).

Mouse embryonic fibroblasts (MEFs) were isolated from E12.5-13.5 embryo by 0.25% Trysin digesting for 10 min and were maintained in complete DMEM media (Sigma, 10% FBS, and 50 U/ml of Penicillin-Stretomycin)

Splenocytes cells were isolated from adult mice by grinding spleen tissue with 40 μM Nilon and culturing in complete RPMI 1640 media (Gibco, 10% HIFBS, 20 mM HEPES, 1 mM sodium pyruvate, 0.05 mM 2-mercaptoethanol, 1X GluMax, and 50 U/ml of Penicillin-Stretomycin). 5 ng/ml of mouse IL-2 was also added into culture media for maintaining splenocytes viability.

### Generation of Crbn knockout and reconstituted Hepa1-6 cell lines

To generate *CRBN* KO cell lines, 293T and Hepa1-6 cells were transfected with the pX459 vector (Addgene, #48139) harboring a specific sgRNA targeting the human *CRBN* locus (5’-GTCCTGCTGATCTCCTTCGC-3’). Following transient transfection, cells were subjected to selection with 1 μg/mL puromycin (Gibco, #A1113803) for 72 h, followed by maintenance in 0.5 μg/mL puromycin for one week. Knockout efficiency was subsequently validated by Western blot analysis.

To establish stable cell lines expressing specific *Crbn* variants, wild-type murine *Crbn* or its mutants (*V380E*, *I391V*, or the *V380E/I391V* double mutant) were cloned into the pLenti6/V5-DEST backbone via homologous recombination. These lentiviral expression vectors were individually co-transfected with the packaging plasmids pCMV-VSV-G and psPAX2 into 293FT cells to produce lentiviral particles.

Supernatants containing the virus were collected 72 h post-transfection, titered, and stored at -80°C. *Crbn* KO Hepa1-6 cells were then infected with the respective lentiviruses overnight. Stable transformants were selected using 5 $\mu$g/mL blasticidin (Life Technologies, #R21001). The resulting cell pools were validated by WB to confirm both the exogenous expression of CRBN variants and the degradation of GSPT1 following CC-885 treatment.

### Generation of *GSPT1^G575N^* knock-in cell lines via RNP electroporation

To generate *GSPT1^G575N^* KI models in 293T and MOLT-4 cells, ribonucleoprotein (RNP) complexes were employed. The RNP complexes, consisting of recombinant Cas9 protein and a synthetic sgRNA targeting the human *GSPT1* locus (5’-TGCTTGGTAGACAAAAAATC-3’), were synthesized by GenScript (Piscataway, NJ, USA). Cells were transfected via electroporation using the Lonza 4D-Nucleofector (program DP-113) with the RNP complexes and 1 μg of single-stranded DNA donor templates harboring the *G575N* mutation.

Following electroporation, cells were cultured in the presence of 500 nM KU-57788 (a DNA-PK inhibitor) for 72 h to enhance the efficiency of homology-directed repair. Single-cell-derived colonies were then isolated through limiting dilution cloning. Genomic DNA from expanded clones was extracted, and the *GSPT1* locus was amplified by PCR and validated via Sanger sequencing. Confirmed *G575N* knock-in clones were further characterized through GSPT1 degradation assays and functional viability tests following CC-885 treatment.

### *In vivo* drug administration

For *in vivo* studies, CC-885 and CC-90009 were initially dissolved in DMSO to create stock solutions. The working solutions were then prepared in a vehicle consisting of 10% DMSO, 40% PEG-300, 5% Tween-80, and 45% sterile saline (0.9% NaCl). Mice were administered the compounds via intraperitoneal injection or oral gavage. The injection volume ranged from 100 to 200 μL per mouse, precisely adjusted according to individual body weight to ensure dosage consistency.

To do the intestine propulsion experiment, active charcoal was spread out evenly in saline with agarose. After 200 ul active charcoal turbid liquid gavage for 30 min, the mouse was sacrificed, and the whole lower gastrointestinal tract (including stomach, intestine, caecum, and colon) was extracted. Using a scissor to cut the mesentery, the intertwining tract could be stretched to measure the length though which activated carbon had moved.

Fresh mouse plasma was obtained by centrifugation of whole blood. Approximately 800 μL to 1 mL of whole blood was collected from the mouse canthus (inner eye corner) and transferred into EDTA-K2 anticoagulant tubes. The tubes were centrifuged at 3000 rpm for 20 min at 4°C. The supernatant (plasma) was then carefully collected, completing the preparation of fresh mouse plasma samples. The plasma samples were maintained on ice and promptly delivered to BIOPHARMA Co., Ltd. for hematological analysis and liver function testing.

### Immunoblot analysis

RIPA buffer was prepared with EDTA-free protease inhibitor cocktail (APExBIO, K1009). The cell pellets were resuspended by RIPA buffer directly to gain whole cells protein lysis, and the protein lysis of various tissues was prepared by grinding the tissues with steel balls in the RIPA buffer at 70hz for 2 min at 4℃. Sonication was optional, but centrifugation was needed for supernate. Protein lysis were quantified with the Pierce BCA Protein Assay Kit (Thermo Fisher). Proteins were separated by SDS-PAGE and transferred onto 0.45 μm pore size Immobilon PVDF membrane (Millipore). PVDF membranes were blocked with TBS-T (50 mM Tris, 150 mM NaCl, and 0.1% Tween-20) containing 5% (w/v) milk. Membranes were incubated with primary antibodies followed by TBS-T washes and incubation with HRP-conjugated secondary antibodies. The signal was visualized with Supersignal West Dura Extended Duration Substrate (Pierce) and ChemiDoc Imaging System (Bio-Rad). The following antibodies were used: anti-GSPT1 (PTM-biolab, PTM6013), anti-CRBN (Sigma, HPA045910), anti-Ikaros (CST, 5443), anti-Aiolos (CST, 15103), anti-ATF4(CST, 11815), anti-Cleaved Caspase3(CST, 9661), anti-Vinculin (Sigma, V9131), and HRP-labeled Goat Anti-Mouse IgG (Epizyme, LF101) or HRP-labeled Goat Anti-Rabbit IgG (Epizyme, LF102).

### Mass spectrum

For cell samples, pellets were harvested by centrifugation at 400 × g for 5 min, washed with ice-cold PBS, and flash-frozen in liquid nitrogen. For tissue samples, specimens were collected, flash-frozen in liquid nitrogen, and stored at -80°C. Tissue samples were further homogenized using steel beads in an ultrasonic cell crusher (Scientz). Both cell and tissue samples were lysed in a buffer containing 8 M urea and 50 mM NH_4_HCO_3_, supplemented with a protease inhibitor cocktail. Following lysis and sonication, the homogenates were centrifuged at 15,000 × g for 12 min at 4°C, and protein concentrations were determined using the Pierce BCA Protein Assay Kit (Thermo Fisher).

For each sample, 150 μg of protein was reduced with 5 mM dithiothreitol (DTT) for 1 h at room temperature (RT) and subsequently alkylated with 10 mM iodoacetamide (IAA) for 30–45 min at RT in the dark. To facilitate enzymatic digestion, samples were diluted with 50 mM NH_4_HCO_3_ to a final urea concentration of 1 M. Protein digestion was performed using sequencing-grade trypsin (Promega) in two steps: an initial digestion at an enzyme-to-substrate ratio of 1:50 at 37°C for 3 h with gentle rotation, followed by the addition of a second equivalent dose of trypsin for overnight incubation at 37°C.

Digested samples were acidified with 1% trifluoroacetic acid (TFA) (Sigma-Aldrich) and desalted on 50 mg tC18 Sep-Pak cartridges (Waters, WAT054960). Cartridges were conditioned with 1 ml 50%vACN/0.1% formic acid (FA), and 1 ml 0.1% FA. Then samples were loaded onto a single cartridge, and subsequently washed twice with 1 ml of 0.1% FA. Samples were eluted from cartridges by washing twice with 1 ml of 50% ACN/0.1% FA. Desalted samples were dried in a Savant SC210A SpeedVac concentrator (Thermo Scientific).

Desalted peptides were separated and analyzed on an Easy-nLC 1200 system coupled to a Q Exactive HF-X (Thermo Scientific). Briefly, about 1 µg of peptides were separated in an Easy-Spray C18 column (75 µm x 50 cm, 2 µm, 100 Å, Thermo Scientific) at a flow rate of 250 nL/min at 50°C. Mobile phase A (0.1% formic acid in water) and mobile phase B (0.1% formic acid in 80% ACN) were used to establish a 90 min gradient. Peptides were then ionized by electrospray at 2.1 kV. A full MS spectrum (375-1500 m/z range) was acquired at a resolution of 120,000 at m/z 200 and a maximum ion accumulation time of 20 ms. Dynamic exclusion was set to 30 s. Resolution for HCD MS/MS spectra was set to 15,000 at m/z 200. The AGC setting of MS and MS^2^ were set at 3E6 and 1E5, respectively. The 20 most intense ions above a 6.7E4 counts threshold were selected for fragmentation by HCD with a maximum ion accumulation time of 30 ms. Isolation width of 1.2 m/z units was used for MS^2^. Single and unassigned charged ions were excluded from MS/MS. For HCD, normalized collision energy was set to 27%. The raw data were processed and searched with MaxQuant 1.6.5.0 with MS tolerance of 4.5 ppm and MS/MS tolerance of 20 ppm. The UniProt mouse protein database and database for proteomics contaminants from Proteome Discoverer were used for database searches. Reversed database searches were used to evaluate t-test of peptide and protein identifications. Two missed cleavage sites of trypsin were allowed. Carbamidomethylation (C) was set as a fixed modification, and Oxidation (M), Acetyl (Protein N-term), and deamidation (NQ) were set as variable modifications. The p-value of both peptide identification and protein identification was set to 0.05. The option of “Second peptides” and “Match between runs” was enabled.

### Immunohistochemistry

Immunohistochemical staining was conducted in paraffin-embedded sections, according to the Mouse and Rabbit Specific HRP/DAB Detection IHC kit (Abcam), with the following antibodies: GSPT1 antibody(1:100) (5) and cleaved Caspase3 antibody (1:200). Sections were counterstained with hematoxylin for 5–10 min, rinsed in running water, differentiated in 1% acid alcohol for 1–3 s, blued in lithium carbonate for 30 s, and washed in distilled water. Slides were dehydrated through graded ethanol (70%, 95%, and 100% for 30 s each), cleared in xylene (2 × 5 min), air-dried, and mounted with neutral resin to avoid air bubbles. Images were captured using a bright-field microscope (Leica). Negative controls were processed in parallel without primary antibody.

### Flow cytometry

Flow cytometry was performed to determine the composition of immune cell subsets in spleen and bone marrow. Freshly isolated spleens and bone marrow cells were gently dissociated and filtered through a 70 μm cell strainer. Cells were washed and resuspended in flow cytometry buffer (PBS containing 2% fetal bovine serum, FBS). After cell counting, equal numbers of cells were aliquoted (100 μL per tube) into 8-well tubes for staining. A multicolor antibody cocktail was prepared in flow buffer. The staining panel included the following antibodies:

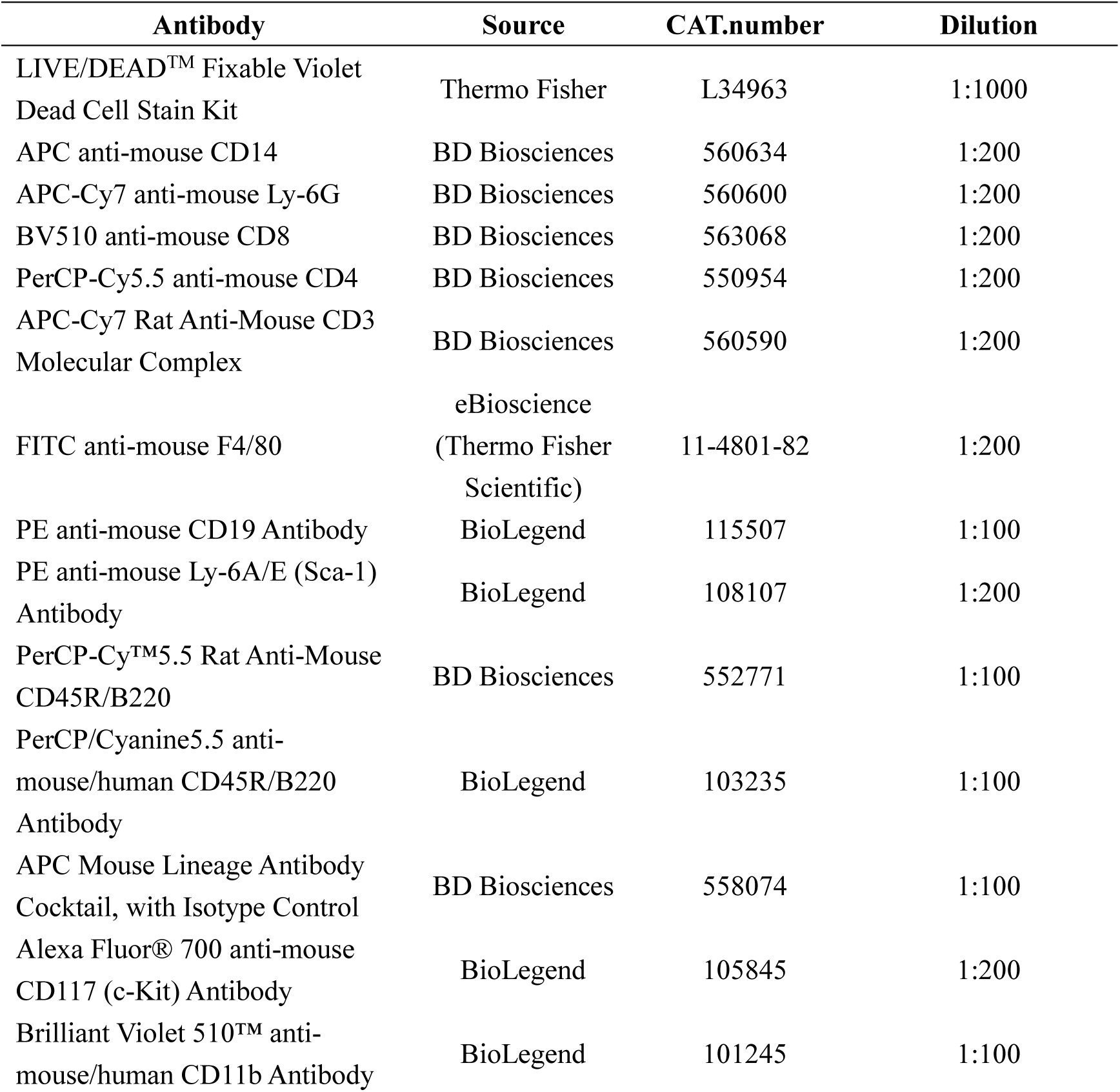
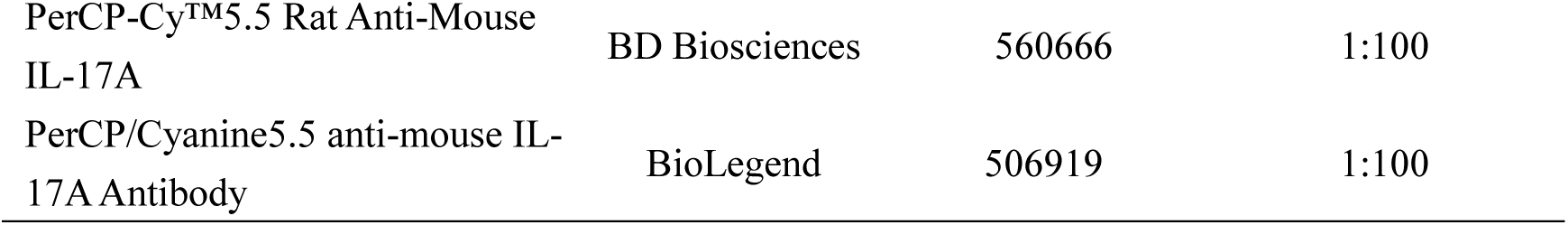

After adding 50 μL of antibody cocktail to each sample, tubes were vortexed briefly and incubated for 30 min at 4°C in the dark. Cells were then washed once with 300 μL flow buffer, filtered through a 40 μm nylon mesh, and resuspended in flow buffer for analysis. Data were acquired on a BD LSRFortessa X20 and analyzed using FlowJo software. Dead cells were excluded using the LIVE/DEAD™ dye. At least 50,000 events per sample were recorded. Cell populations were identified using the following gating strategies:

| Subset | Gating markers |
| --- | --- |
| Monocytes | CD45<sup>+</sup> CD11b<sup>+</sup> F4/80<sup>-</sup> CD14<sup>+</sup> |
| Neutrophils | CD45<sup>+</sup> CD11b<sup>+</sup> F4/80<sup>+</sup> Ly-6G<sup>+</sup> |
| CD4<sup>+</sup> T cells | CD45<sup>+</sup> CD3<sup>+</sup> CD4<sup>+</sup> |
| CD8<sup>+</sup> T cells | CD45<sup>+</sup> CD3<sup>+</sup> CD8<sup>+</sup> |
| B cells (spleen) | CD45<sup>+</sup> CD19<sup>+</sup> |
| Hematopoietic stem cells (HSCs) | Lin<sup>-</sup> Sca-1<sup>+</sup> CD117<sup>+</sup> |
| Myeloid cells | CD11b<sup>+</sup> |
| Total B cells (bone marrow) | B220<sup>+</sup> CD19<sup>+</sup> |

### RNA sequencing and analysis

The liver and spleen were immediately harvested following mouse euthanasia. Each organ was divided into three to four aliquots and flash-frozen in liquid nitrogen for preservation. For 293T cell samples, trypsin-digested cells were collected into 1.5 mL microcentrifuge tubes and similarly flash-frozen in liquid nitrogen. Samples were sent to Novogene Co., Ltd. for RNA extraction, library construction, and sequencing. RNeasy MinElute Cleanup Kit (QIAGEN, 74204) was used for RNA extraction, followed by generating mRNA-focused sequencing libraries from total RNA using Illumina® TruSeq® RNA Sample Preparation Kit v2. Paired-end 150 bp sequencing was done on illumina Hiseq X10 (40 M reads for each sample).

The transcriptomic analysis work flow began with a thorough quality check by FastQC (0.11.2) and the adapters of each sample were trimmed respectively. The raw reads were mapped to the mouse (Mus musculus) genome (mm10) using STAR36 (2.4.2a) and annotated with transcriptome database (genecode vM13). Gene abundance estimation by reads count was conducted with the software RSEM37 (1.2.29). Normalized tags per million (TPM) were calculated on the number of clean reads mapped to specific region of genome using the relative log expression (RLE) method in edgeR38 (3.16.5). Differentially expressed genes (DEG) refer to compare gene expression level between two samples or two groups. DEG were defined by using the criterion that fold change >1.6 and adjust P < 0.05 (adjust P-value). We employed GSEA to explore the DEG and TPM. To perform GSEA, we utilized gene sets available on the Molecular Signatures Database website (https://www.gsea-msigdb.org/). Normalized Enrichment scores (NES) and false discovery rates (FDR) are denoted within the figures.

### Real-Time PCR

Total RNA was extracted from the spleen with Trizol reagent. cDNAs were synthesized from 1 µg of total RNA using the SPARKscritpt II RT Plus Kit (SparkJade) and were amplified by 2×SYBR Green qPCR Mix (SparkJade) using Quantstudio 7 Real-Time PCR System (Life Technologies) according to the manufacturer’s protocols. The following primers were used for qPCR:

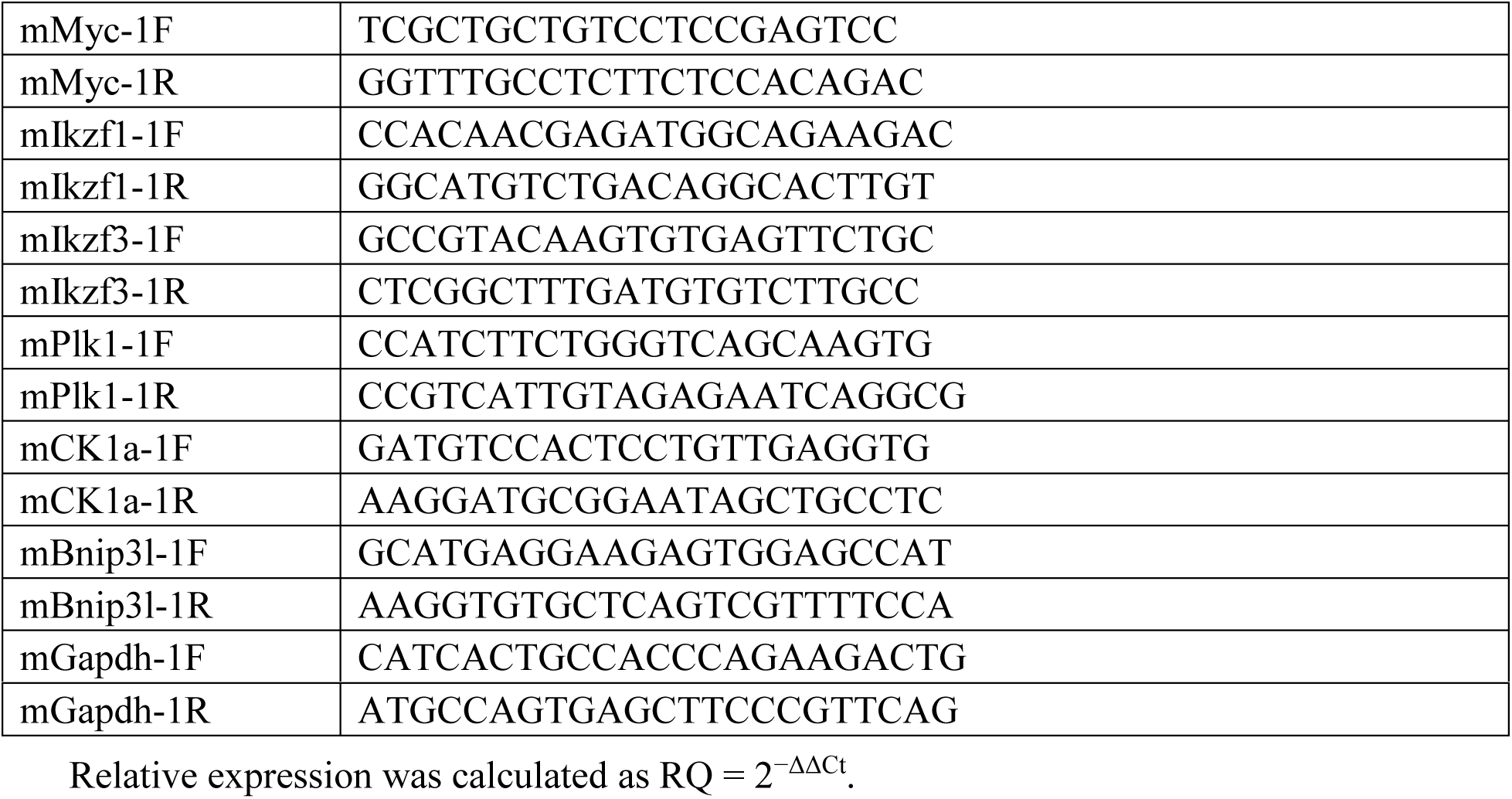

### Statistical analysis

For all experiments, the number of technical and/or biological replicates is provided in the figure legends or text. All statistical analyses were performed using GraphPad Prism 8 software. Comparisons between two treatments were assessed using unpaired or paired (for matched comparisons) two-tailed Student’s t-test. In all cases, n.s. no significance.

## Supplemental information

**Figure S1.**
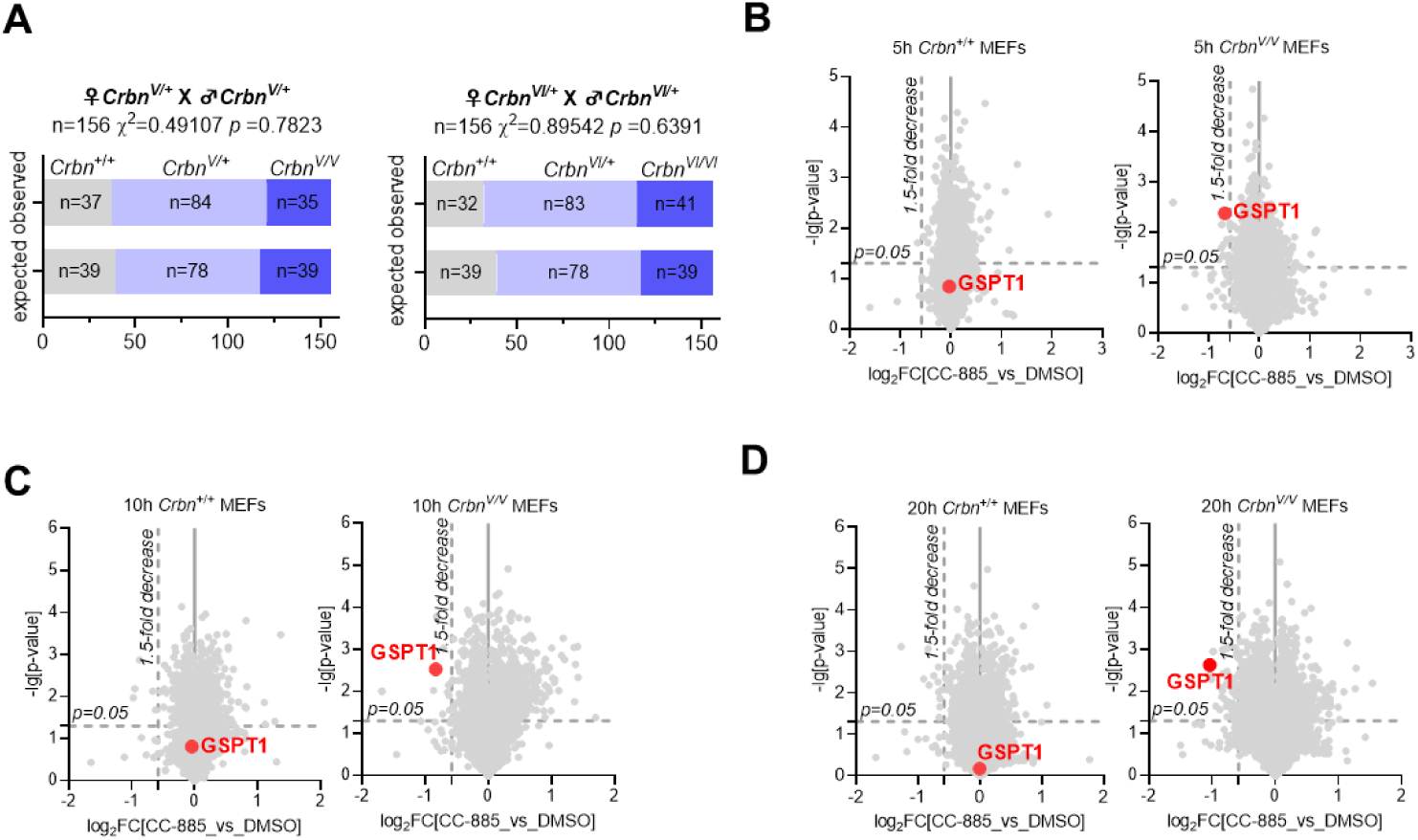
Generation and validation of *Crbn^V/V^* and *Crbn^VI/VI^* mouse models. (A) Genotype distribution of offspring from *Crbn^V/+^*and *Crbn^VI/+^* heterozygous intercrosses. Observed frequencies of F1 genotypes were recorded, with Chi-square tests performed to assess adherence to Mendelian inheritance patterns. (B-D) Quantitative proteomic analysis of GSPT1 degradation and global protein profiles. *Crbn*^+/+^ and *Crbn*^V/V^ MEFs were treated with CC-885 for the indicated durations (5, 10, and 20 h). Protein levels were quantified via TMT-labeled mass spectrometry to evaluate the kinetics of degradation and proteome-wide selectivity.

**Figure S2.**
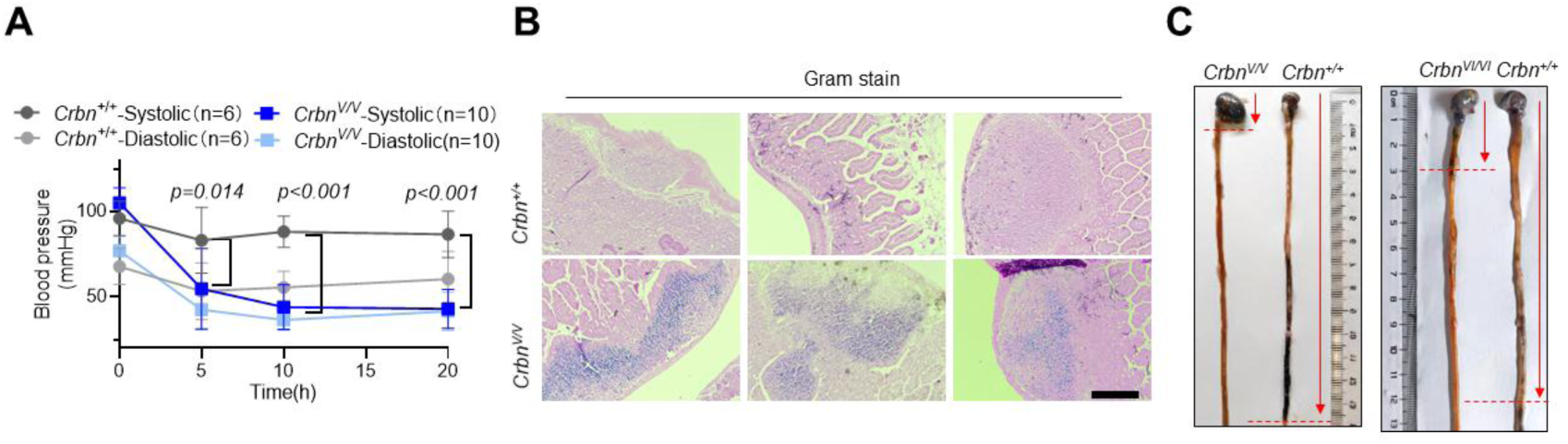
GSPT1 degraders induce intestinal toxicity in *Crbn* humanized mice. (A) Physiological monitoring of blood pressure. Systolic and diastolic blood pressure were measured via a tail-cuff system in *Crbn*^+/+^ and *Crbn^V/V^*mice following a single intraperitoneal injection of 5 mg/kg CC-885. Data are presented as mean ± SEM (*n* ≥3 biological replicates). *P* values were determined by multiple t-tests. (B) Representative images of Gram-stained intestinal sections. Gram-positive bacteria (blue-violet) were visualized in the intestinal tissues of *Crbn*^+/+^ or *Crbn^V/V^* mice 20 h post-treatment with 5 mg/kg CC-885 (single *i.p.* dose). Scale bar, 200 μm. (C) Assessment of gastrointestinal motility. Intestinal transit was evaluated 15 h after a single *i.p.* administration of 5 mg/kg CC-885 using a charcoal propulsion assay in *Crbn*^+/+^ and *Crbn^V/V^* mice.

**Figure S3.**
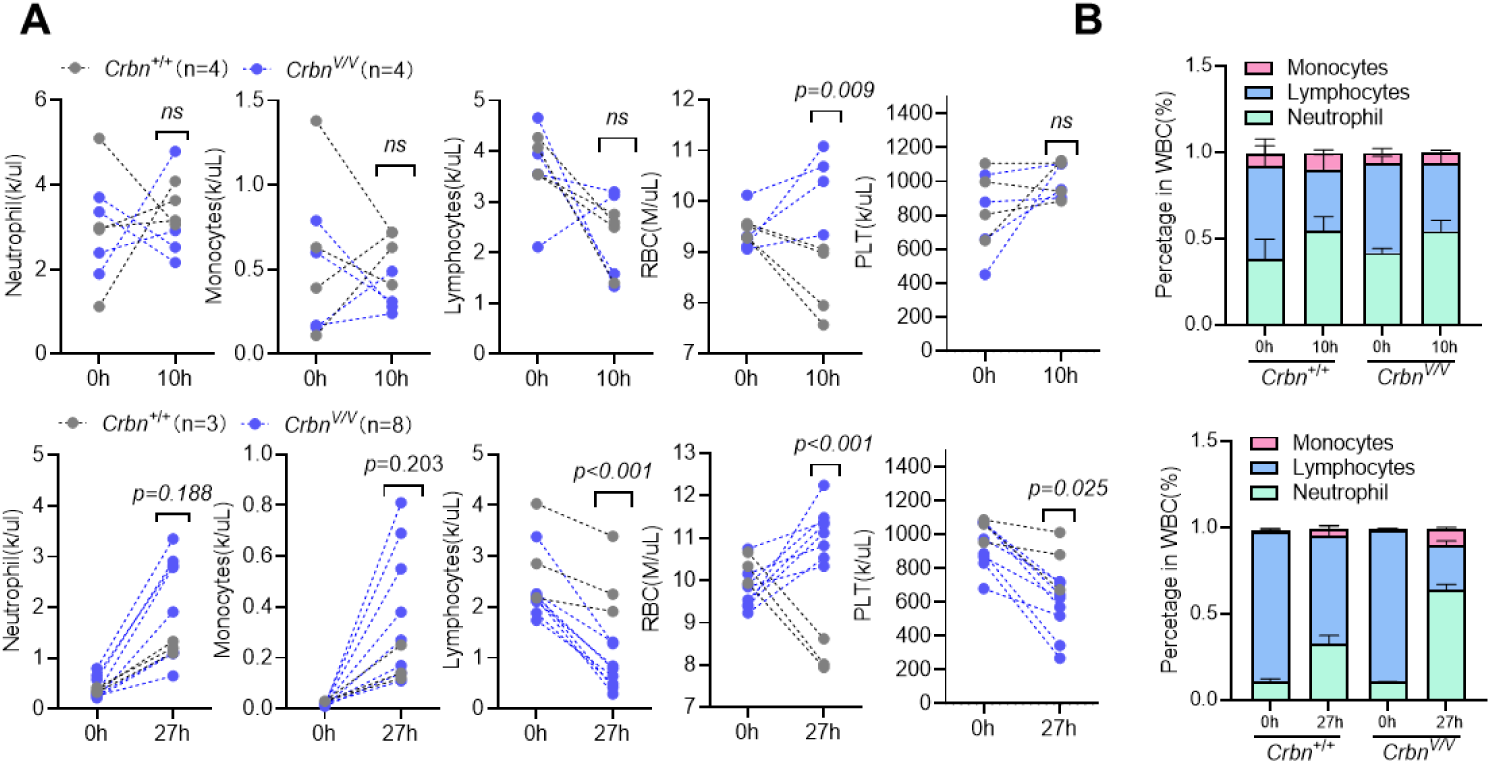
CC-885 induces hematological alterations in *Crbn^V/V^* mice, characterized by erythrocytosis, leukopenia, neutrophilia, and monocytosis. (A) Absolute hematological counts. Absolute counts of RBCs, leukocyte subsets (three-part differential), and platelets in *Crbn^V/V^* mice at 10 h (top) and 27 h (bottom) post-injection of 5 mg/kg CC-885. Data are presented as mean ±SEM (*n* ≥3 biological replicates). *P* values were determined by multiple t-tests. (B) Leukocyte differential percentages. Relative percentages of monocytes, lymphocytes, and neutrophils within total leukocytes at 10 h and 27 h post-injection.

**Figure S4.**
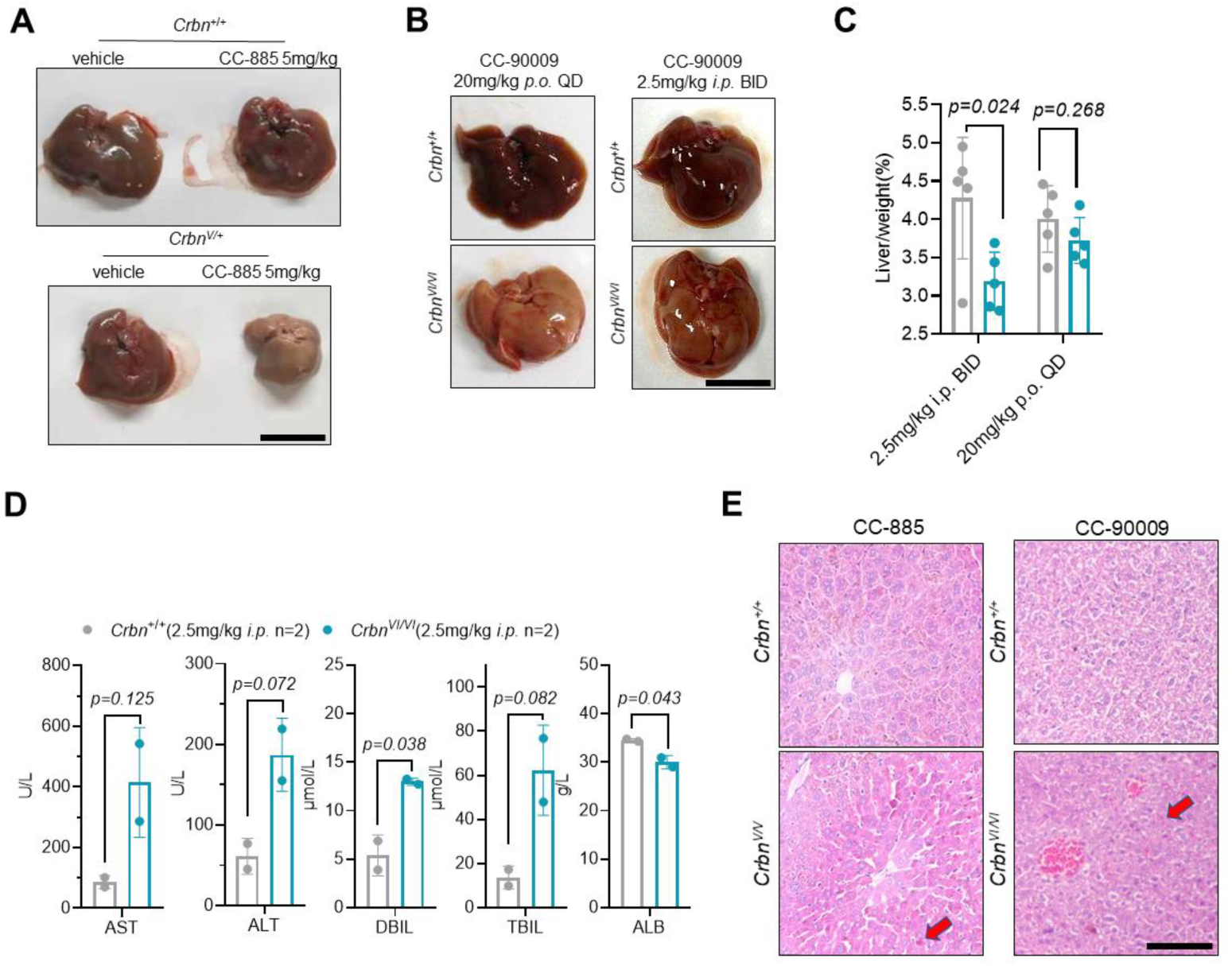
GSPT1 degraders induce hepatic congestion and functional impairment in humanized *Crbn* mice. (A) Macroscopic liver morphology (CC-885). Dissection microscopy of livers from *Crbn*^+/+^ and *Crbn*^V/+^ mice 20 h post-administration of 5 mg/kg CC-885 (single *i.p.* dose). Scale bar, 10 mm. (B) Macroscopic liver morphology (CC-90009). Dissection microscopy of livers from *Crbn*^+/+^ and *Crbn*^VI/VI^ mice following CC-90009 treatment at 20 mg/kg (*p.o.*, QD) or 2.5 mg/kg (*i.p.*, BID). Scale bar, 10 mm. (C) Liver weight analysis. Distribution of liver weights from the mice described in **Figure S4B**. (D) Hepatic functional. Plasma liver biomarkers in *Crbn*^+/+^ and *Crbn*^VI/VI^ mice following 2.5 mg/kg CC-90009 (*i.p.*, BID). For the treatment group in **Figure S4C**, only 2 out of 5 mice survived to the final time point for plasma collection. (E) Hepatic histological assessment. Corresponding liver sections show congestion and stasis (red arrowheads). Scale bar, 100 μm. Data: mean ±SD (**C**); *n*, biological replicates. *P* values: multiple t-tests. All experiments performed more than 3 times with representative data shown. n.s., not significant.

**Figure S5.**
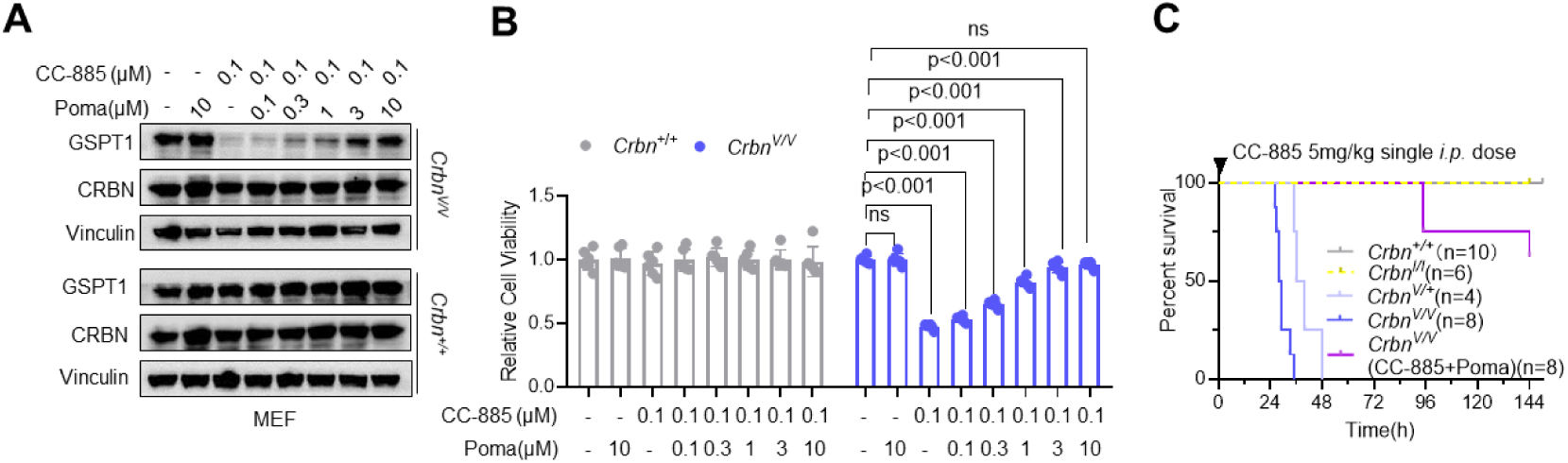
Competitive inhibition of Crbn by pomalidomide rescues GSPT1 degradation and prolongs survival in *Crbn^V/V^* mice. (A) Rescue of GSPT1 degradation. Immunoblot analysis of GSPT1 levels in *Crbn*^+/+^ and *Crbn^V/V^* MEFs following 12 h co-treatment with Poma and CC-885. (B) Cell viability. CCK-8 assay of *Crbn*^+/+^ and *Crbn^V/V^* MEFs after 48 h co-treatment with Poma and CC-885. Data are presented as mean ±SEM (*n* ≥6 biological replicates). *P* values were determined by multiple t-tests. (C) In vivo survival. Survival curves of *Crbn*^+/+^ and *Crbn^V/V^* mice treated with 5 mg/kg CC-885 with or without 40 mg/kg Poma.

**Figure S6.**
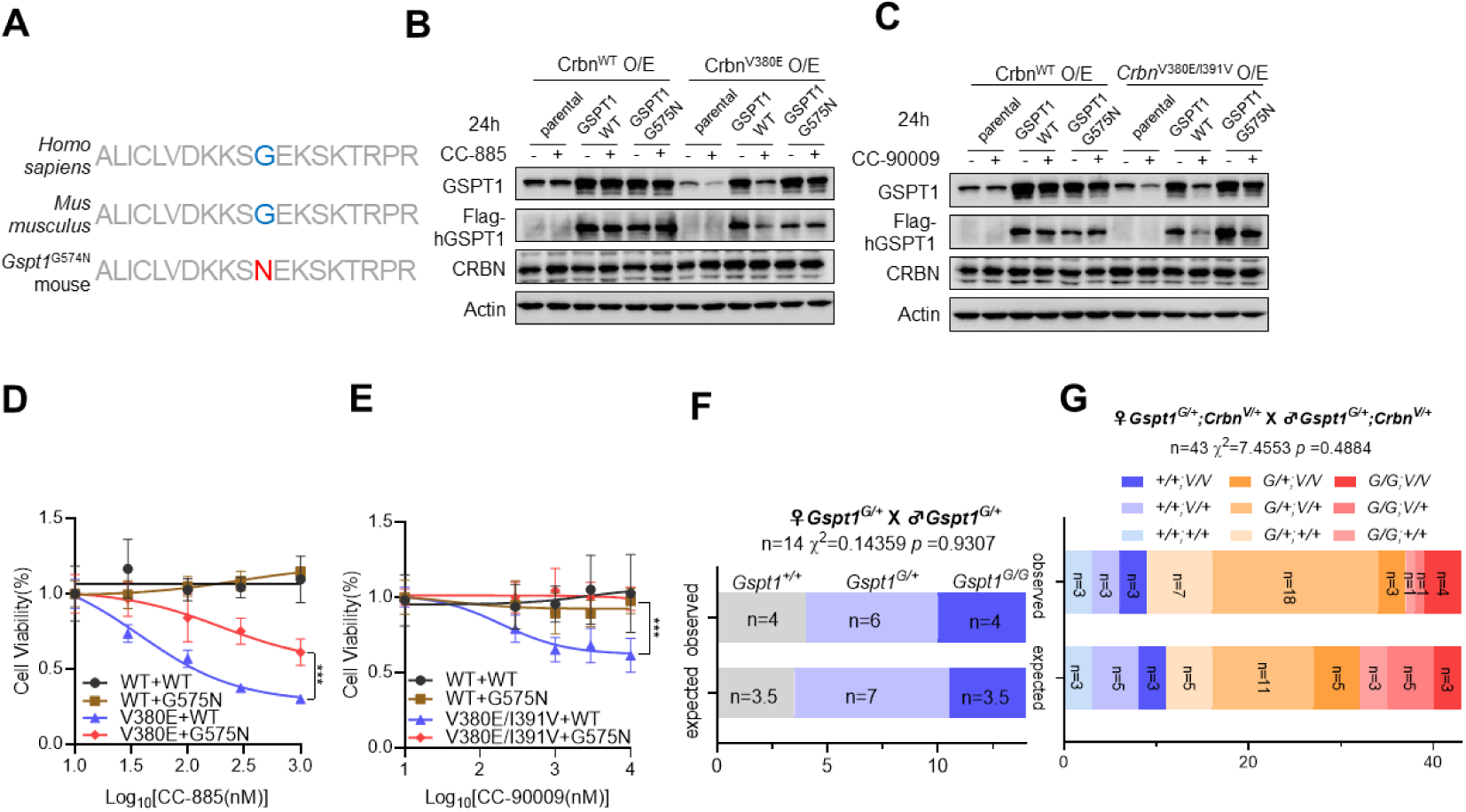
Generation and validation of the degrader-resistant *Gspt1^G574N^* mouse model. (A) Sequence homology. Sequence alignment of the G-loop region in GSPT1 between human and mouse, highlighting the conserved glycine at positions 575 (human) and 574 (mouse). (B and C) Functional validation in mouse *Crbn* KO cells. Immunoblot analysis of GSPT1 degradation in *Crbn* KO Hep1-6 cells co-transfected with either WT or humanized mouse Crbn, along with WT GSPT1 or the resistant GSPT1^G575N^ mutant. Cells were treated with 0.1 μM CC-885 (B) or 1 μM CC-90009 (C) for 24 h. (D and E) Cell viability. Growth inhibitory effects of CC-885 (D) and CC-90009 (E) on the Hep1-6 cells described in (B) and (C) after 48 h of treatment. (F and G) Mendelian inheritance analysis. Genotype distribution of offspring from heterozygous intercrosses for *Gspt1^G574N^*single-mutant (F) and *Crbn^V380E^*; *Gspt1^G574N^* double-mutant (G) strains. Observed frequencies were compared against expected Mendelian ratios using the chi-square test.

**Figure S7.**
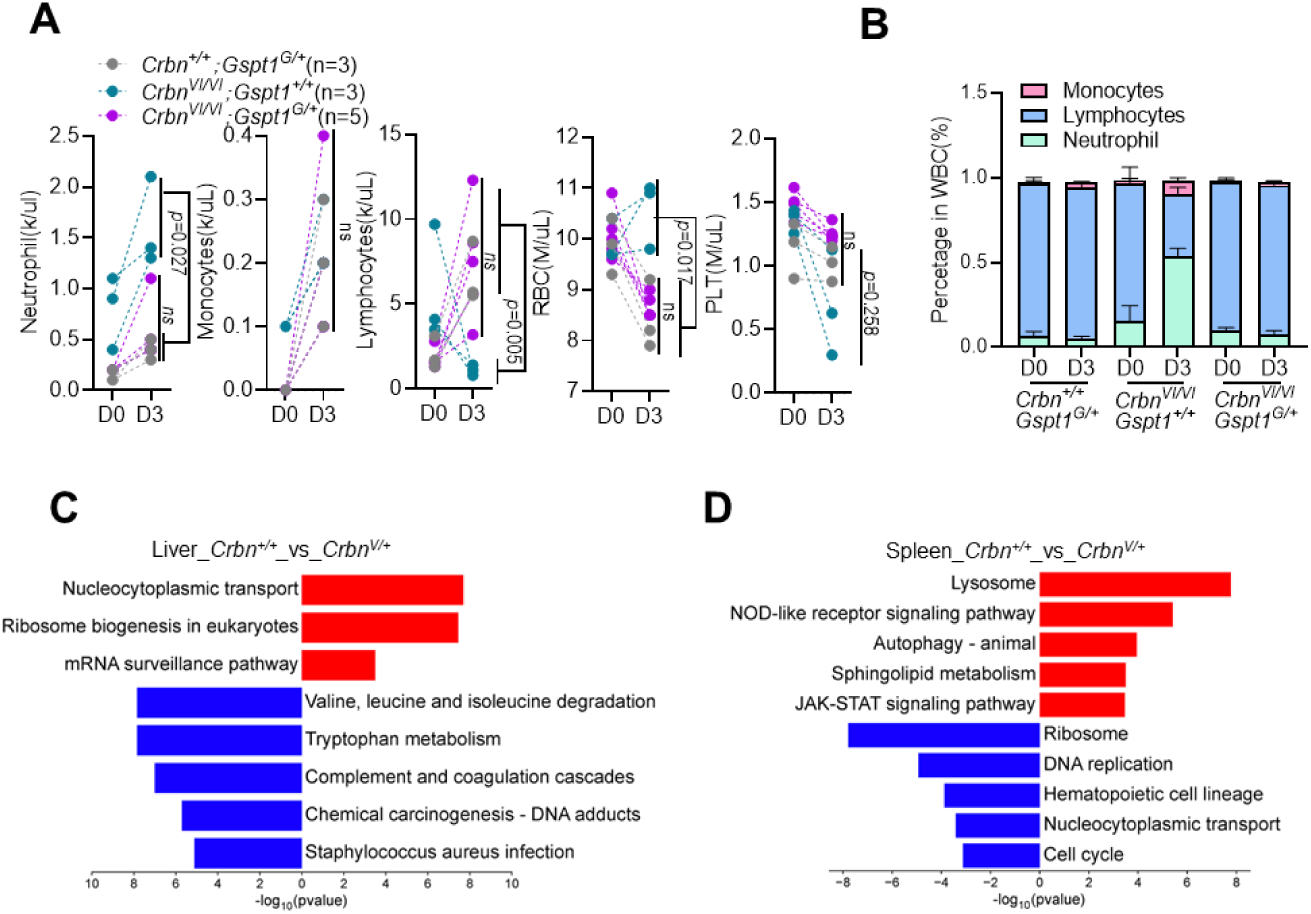
The *Gspt1^G574N^* mutation confers resistance to multi-organ damage in humanized Crbn mice. (A and B) Hematological profiles. Absolute blood cell counts (A) and relative leukocyte proportions (B) in *Crbn*^+/+^, *Crbn^VI/VI^* and *Crbn^VI/VI^*; *Gspt1^G/+^*mice following a 3-day treatment regimen of CC-90009 (20 mg/kg, *i.p.*, QD). (C and D) Transcriptomic signatures. GSEA (Gene Set Enrichment Analysis) of enriched biological pathways in the liver (C) and spleen (D) from the experimental groups described in Figure 4A and 4B.

**Figure S8.**
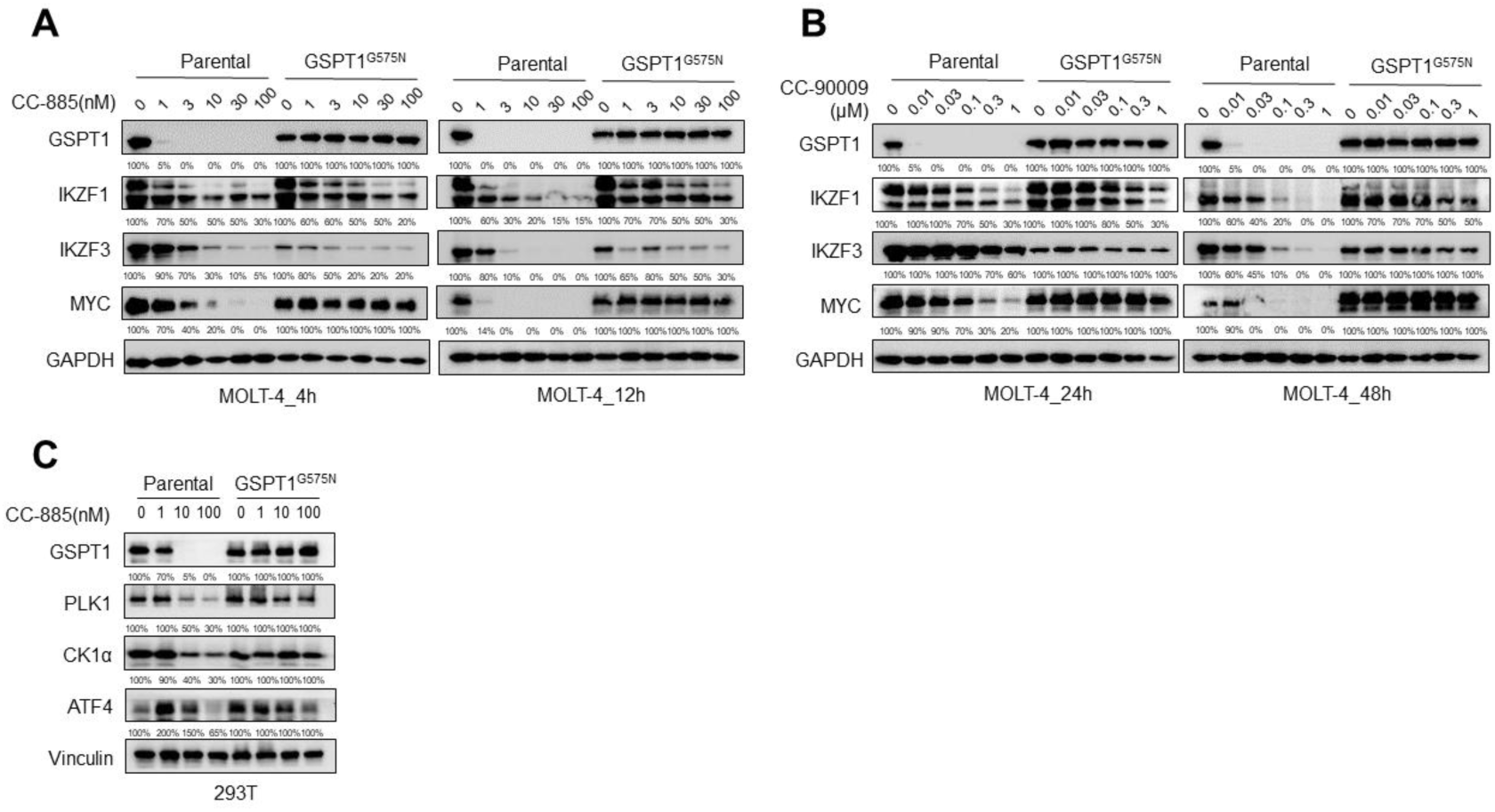
GSPT1 degradation leads to secondary alterations in downstream protein abundance in human cells. (A and B) Comparative protein expression in MOLT-4 cells. Immunoblot analysis of GSPT1, IKZF1/3, and MYC in WT and *GSPT1^G575N^* mutant MOLT-4 cells. Cells were treated with gradient concentrations of CC-885 (A) or CC-90009 (B) for the indicated durations. (C) Protein levels in 293T cells. Western blot analysis of GSPT1, PLK1, CK1α, and ATF4 in WT and *GSPT1^G575N^* mutant 293T cells following 12 h treatment with gradient concentrations of CC-885.

**Figure S9.**
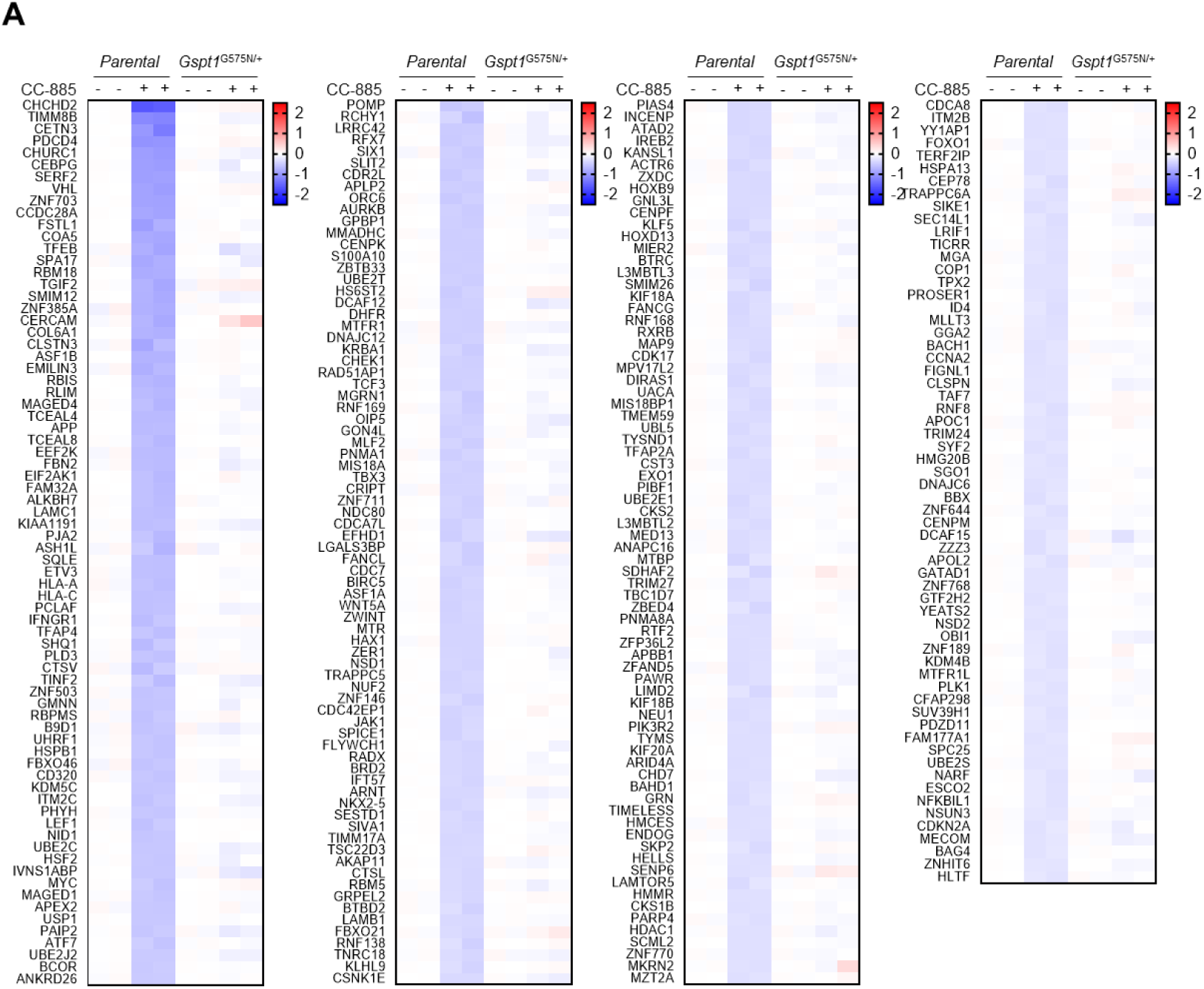
Analysis of the downstream proteomic landscape following GSPT1 degradation. (A) Representative list and heatmap (as described in Figure 5) of candidate proteins exhibiting decreased abundance as a secondary consequence of GSPT1 degradation. This signature includes short-lived proteins and other translationally sensitive targets whose homeostatic levels are disrupted following the loss of GSPT1-mediated translation termination.

**Figure S10.**
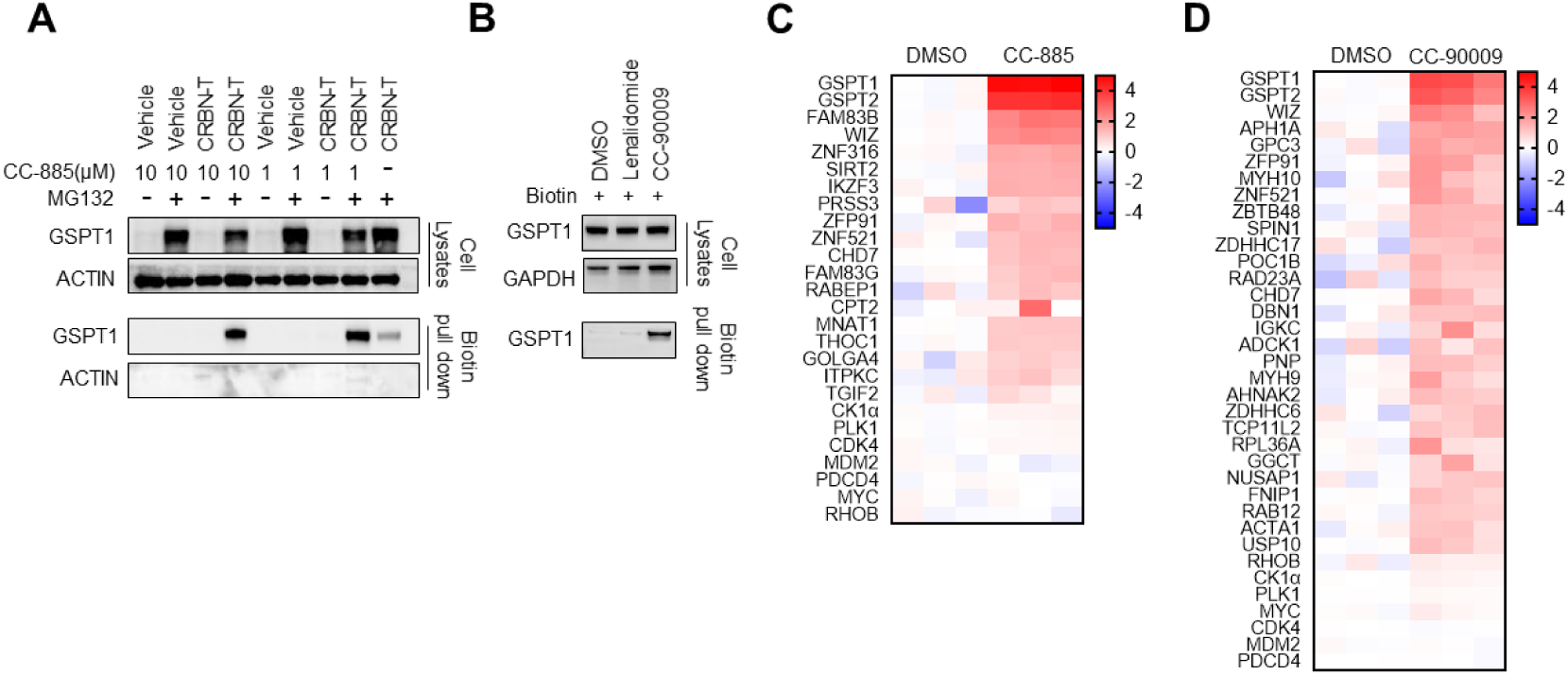
Identification of proteins recruited by CRBN in the presence of GSPT1 degraders. (A and B) Validation of the TurboID system. Immunoblot analysis of GSPT1 recruitment in CRBN-TurboID expressing 293T cells treated with CC-885 (A) or CC-90009 (B). The enrichment of GSPT1 in the pull-down fraction confirms the functional integrity of the proximity-labeling system. (C and D) Candidate protein recruitment profiles. Heatmaps showing selected proteins identified by mass spectrometry following CRBN-TurboID labeling with CC-885 (C) or CC-90009 (D). The color intensity (red) represents the relative probability of recruitment to the CRBN complex.

**Table S1.**
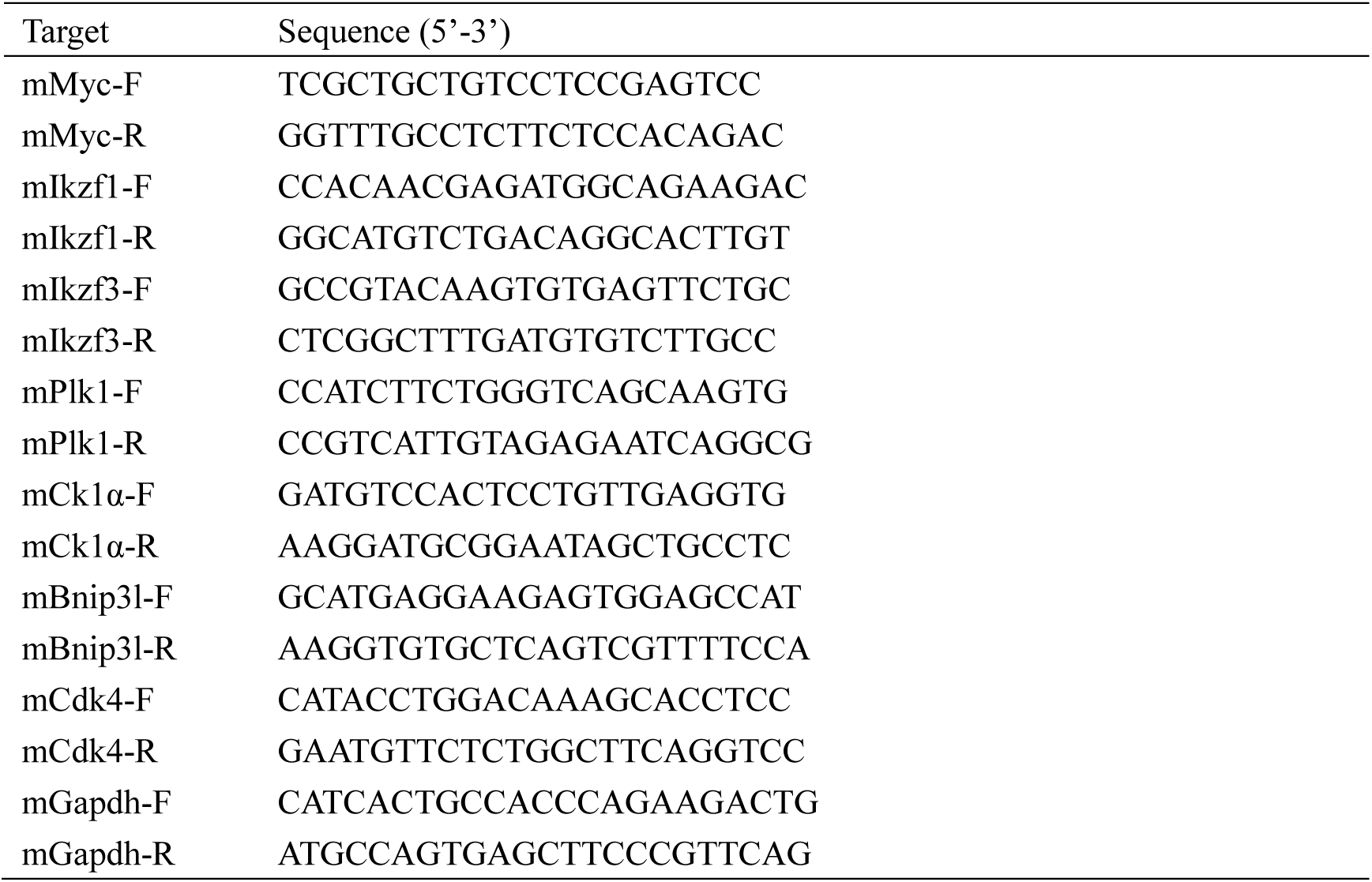
Primers for Real-Time PCR. Related to Figure 4.

